# Breast cancer identity is defined by specialized enhancer sets via lysine deacetylation

**DOI:** 10.1101/2025.07.31.668008

**Authors:** Hiroaki Tachiwana, Kohei Kumegawa, Ryota Matsudo, Ai Katsuma, Atsushi Okabe, Nao Yoshida, Xufeng Shu, Masaki Kato, Tamiko Minamisawa, Akihiro Ito, Hiroshi Kimura, Piero Carninci, Atsushi Kaneda, Yasukazu Daigaku, Reo Maruyama, Noriko Saitoh

## Abstract

Breast cancer subtypes are defined by distinct transcriptional programs, yet the epigenetic mechanisms underlying subtype-specific gene regulation remain unclear. Enhancers, key regulators of gene expression and cell identity, are well positioned to define breast cancer subtypes. Here, we identify a previously unrecognized class of enhancers, termed hypoacetylation-defined (HD) enhancers, that regulate cancer-related genes in a luminal breast cancer cell line. HD enhancers are defined by RNA polymerase II dissociation upon lysine deacetylase inhibition, and bidirectional eRNA transcription. They are distinct from super-enhancers, require a specific Mediator subunit for gene-specific transcription, and form extensive chromatin interactions suggestive of a hub-like architecture. Analyses of clinical datasets further identified a subset of HD enhancers, termed HD cluster 1 enhancers, which classify patients into breast cancer subtypes and are associated with expression quantitative trait loci linked to subtype-specific gene expression. This study identifies the lysine deacetylation-regulated cell identity enhancers, which are potential therapeutic targets.

## Introduction

Breast cancer is a biologically and clinically heterogeneous disease^1^. A set of 50 genes, known as PAM50, was proposed to accurately classify breast cancer into molecular intrinsic subtypes: luminal A, luminal B, HER2-enriched, basal-like and normal-like subtypes^2^. Recent studies have highlighted the importance of epigenetic regulation in shaping these subtype-specific transcriptional programs, particularly through chromatin-based mechanisms^3^. Enhancers, cis-regulatory elements that modulate gene expression through physical interactions with their target promoters, have emerged as key determinants of cell identity and are likely to contribute to subtype-specific gene regulation in breast cancer. Supporting this, recent analyses have shown that the risk-associated noncoding variants identified by genome-wide association studies (GWAS) tend to reside in putative enhancer regions^4,5^. Additionally, expression quantitative trait loci (eQTL), which have genetic variants that affect gene expression, are often located within enhancers^6,7^.

Enhancers are marked by specific histone modifications, such as histone H3 monomethylation at lysine 4 (H3K4me1) and acetylation at lysine 27 (H3K27ac)^8–10^, and are often found in DNase I hypersensitive regions, reflecting their open and accessible chromatin architecture^11,12^. Based on these features, putative enhancers have been identified throughout the human genome, which is estimated to contain nearly one million enhancers across different cell types^9,13^. Enhancers are also identified through the bidirectional transcription of non-coding enhancer RNAs (eRNAs) by RNA polymerase II (RNAPII)^14^, which can be mapped by cap analysis of gene expression (CAGE) sequencing and serves as a reliable marker of active enhancers^15–17^.

Alternatively, genome-wide enhancer mapping has been achieved through the integration of chromatin accessibility (ATAC-seq) and histone modification (ChIP-seq) data, leading to the identification of candidate cis-regulatory elements (cCREs), a subset of which are putative enhancers^18^. Although chromatin accessibility alone does not define enhancer function, distal regions showing cell state-specific accessibility are frequently enriched for enhancer-associated features and have been shown to drive gene expression *in vivo*^19,20^. Accordingly, ATAC-seq data can serve as a useful proxy for identifying candidate enhancer regions across the genome. In addition, enhancer-promoter interactions can be analyzed at high resolution genome-wide using chromatin conformation capture techniques such as 4C and Hi-C^21–24^, enabling the functional linkage of distal enhancers to their target genes.

Moreover, enhancers vary in their molecular composition. The Mediator complex functions as a critical co-activator in transcription by bridging enhancers and promoters^25–27^. Its subunit composition varies across target genes, and inhibition of specific subunits results in gene-specific transcriptional outcomes^28,29^. This structural and functional diversity allows the Mediator complex to fine-tune transcriptional programs in a context-dependent manner, acting through diverse classes of enhancers to shape the gene expression patterns that define cell identity. Among these enhancer classes, super-enhancers, regions with exceptionally high densities of transcription factors and Mediator binding, have been linked to genes that regulate cell identity^30^. However, most enhancers fall outside this category and may operate through distinct and incompletely understood regulatory logic.

Deacetylation of histone and non-histone proteins may contribute to the regulation of transcriptional programs that define cell identity. Histone deacetylases (HDACs) are generally linked to transcriptional repression, although they have also been reported to function at actively transcribed loci^31^. Furthermore, the inhibition of HDACs reduces the expression of specific genes, including several oncogenes in tumor cells^32,33^. HDAC activity is also required for maintaining pluripotency gene expression through the recruitment of RNAPII^34^. Since HDACs also act on non- histone proteins^35,36^, these findings suggest that the HDAC-mediated deacetylation of histone and/or non-histone proteins may help shape the cell identity-defining transcriptomes. Nevertheless, the precise mechanisms by which HDAC activity contributes to the maintenance of cell identity-defining transcriptional programs remain poorly understood.

In this study, we have identified a previously unrecognized class of enhancers, termed hypoacetylation-defined (HD) enhancers, that support the cell identity of luminal breast cancer MCF-7 cells. Their activity is disrupted upon histone deacetylase (HDAC) inhibition, accompanied by the reduced recruitment of RNAPII and the Mediator complex, leading to the repression of key cell identity genes. An integrative analysis of patient-derived datasets further revealed a subset of 216 HD Class 1 (HDC1) enhancers whose chromatin accessibility stratifies with PAM50-defined intrinsic subtypes. These enhancers are also enriched for eQTLs associated with luminal-specific gene expression programs. Our findings highlight a distinct enhancer logic that operates in the presence of HDAC activity and plays a critical role in sustaining the luminal cancer-specific transcriptional networks. This enhancer class not only provides insight into the epigenetic architecture of cancer cell identity but also presents promising avenues for subtype-specific biomarkers and therapeutic intervention strategies beyond conventional enhancer models.

## Results

### Genes associated with breast cancer properties are down-regulated upon hyperacetylation

To investigate whether lysine deacetylation contributes to shaping cell identity-defining transcriptomes in breast cancer cells, we performed RNA-seq analysis of the human luminal breast cancer cell line MCF-7 after a 3 h treatment with SAHA (suberoylanilide hydroxamic acid), an inhibitor of multiple HDACs. The short treatment duration was designed to be short, to capture the direct transcriptional effects of SAHA and minimize secondary responses^34,37,38^. Western blot and ChIP-seq data showed that the histone acetylation levels increased as expected (Figures 1A and 1B). Based on the RNA-seq data, we identified differentially expressed genes, including 212 SAHA-up-regulated and 330 SAHA-down-regulated genes, as well as 2,564 SAHA-unchanged genes (Figure 1C). Representative genome browser tracks supported these classifications (Figure 1D).

**Figure 1.**
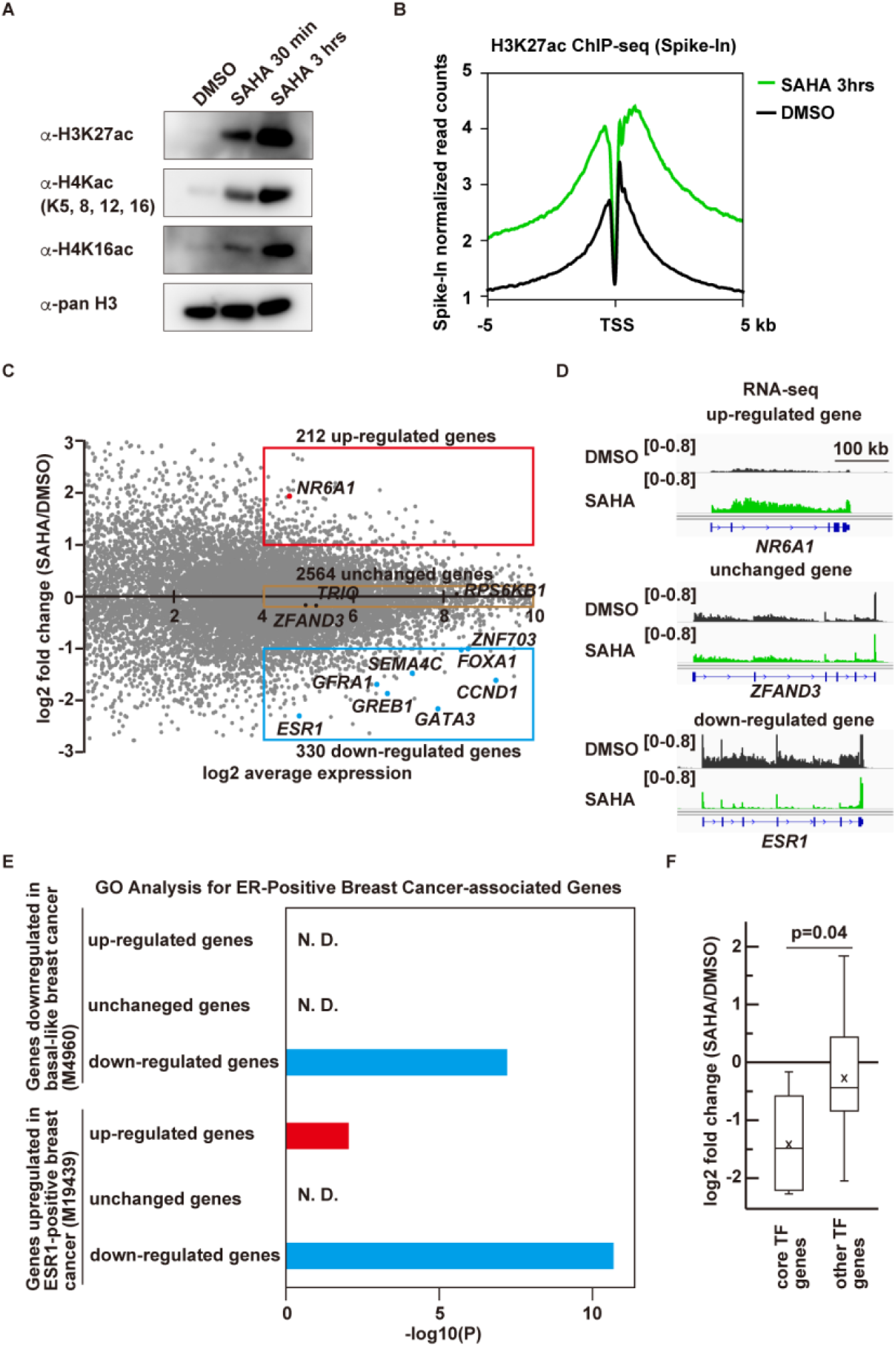
The deacetylase inhibitor SAHA repressed cell identity gene transcription in MCF-7 cells. (A) The luminal breast cancer MCF-7 cells were treated with 2 µM SAHA or DMSO for 30 min or 3 h, and acetylated histone H3 and H4 were examined with western blots. Pan H3 served as a loading control. (B) Aggregation plot of H3K27ac ChIP-seq data at transcription start sites (TSS) in cells with (green) and without (black) 3 h SAHA treatment. SAHA increased the acetylation levels in chromatin. The data were normalized using spike-in controls. (C) The MA plot of total RNA-seq of cells after 3 h SAHA treatment, showing up-regulated (red; n = 212, log2 fold change > 1), down-regulated (blue; n = 330, log2 fold change < -1), and unchanged (brown; n = 2,575, log2 fold change between -0.2 and 0.2) genes, with log2 average expression > 4. (D) RNA-seq tracks of up-regulated (*NR6A1*), unchanged (*ZFAND3*), and down-regulated (*ESR1*) genes. RNA-seq tracks for DMSO (black) and SAHA (green) treatments are aligned from left to right, indicating transcription direction. (E) GO analysis using MSigDB (GSEA), showing that SAHA-down-regulated genes are enriched in luminal breast cancer-related genes (blue bars). (F) Core TF genes were down-regulated following the SAHA treatment. Core TF genes include *ESR1*, *GATA3*, *FOXA1*, *SPDEF*, and *TFAP2C*, while other TF genes include 116 genes with a log2 average expression > 4.

A GO analysis of the three groups confirmed that the genes repressed by HDAC are enriched in the SAHA-upregulated genes (Figure S1). Interestingly, the SAHA-down-regulated group included genes associated with luminal breast cancer, including *ESR1*, *GATA3*, *GREB1*, and *CCND1* (Figures 1C, 1E and S1). This group also included five core transcription factor (core TFs) genes of MCF-7^39^, *ESR1*, *GATA3*, *FOXA1*, *TFAP2C*, and *SPDEF*, which were significantly more down-regulated than other TFs (Figure 1F). Thus, SAHA-induced hyperacetylation repressed a specific set of luminal breast cancer-related genes.

### Genes with high RNAPII binding are susceptible to transcriptional repression by SAHA

To investigate SAHA-induced transcriptional repression, we examined histone acetylation near transcription start sites (TSSs) by ChIP-seq. The H3K27 acetylation increased to similar levels in down-regulated, unchanged and up-regulated genes upon the SAHA treatment (Figure 2A). These results suggest that SAHA represses the transcription of specific genes despite increased histone acetylation, possibly via the acetylation of non-histone proteins.

**Figure 2.**
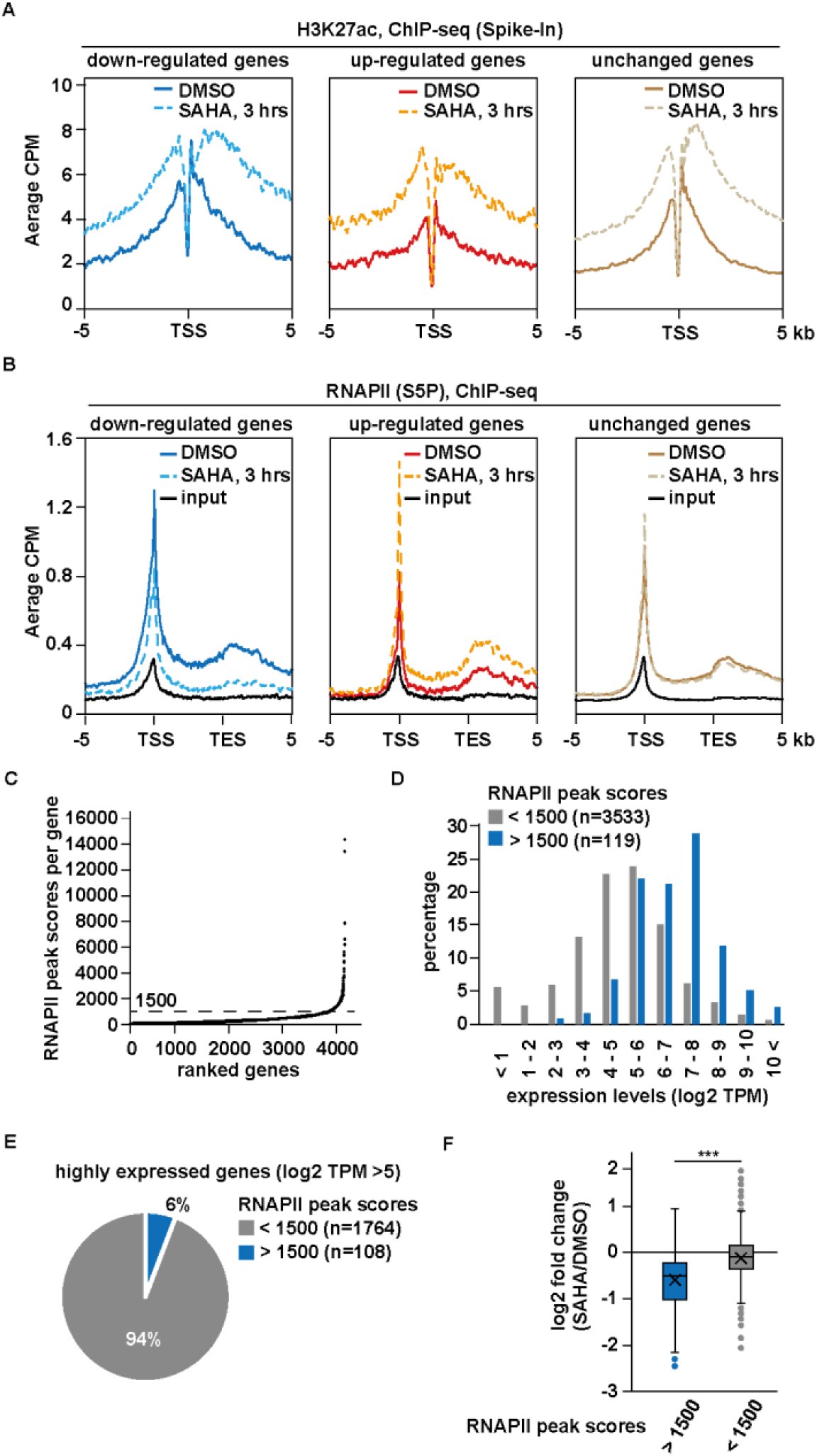
RNAPII is densely bound to SAHA-down-regulated genes and dissociated upon SAHA treatment. (A) ChIP-seq profiling of H3K27ac upon SAHA treatment (dashed line) and DMSO control (solid line). H3K27ac levels increased in all gene groups, including SAHA-down-regulated (left), up- regulated (middle), and unchanged (right) genes. ChIP-seq data were normalized using spike-in controls. (B) ChIP-seq profiling of transcriptionally active RNAPII (S5P). Upon SAHA treatment, RNAPII association decreased in SAHA-downregulated genes (left), increased in SAHA-upregulated genes (middle), and remained unchanged in SAHA-unchanged genes (right). Under basal conditions (DMSO), SAHA-down-regulated genes exhibit higher RNAPII binding levels, as compared to other genes. The input control for RNAPII ChIP-seq is represented by a black line. (C) RNAPII (S2P) binding levels for each gene were ranked based on the ChIP-seq data of MCF-7 cells. RNAPII peak scores per gene were calculated by summing all peaks. (D) Distributions of expressed genes with high RNAPII binding (blue bars, peak score > 1,500) and other genes (gray bars), showing that genes with high RNAPII binding have high expression levels. (E) Genes densely bound by RNAPII represent only 6% of highly expressed genes (log2 TPM > 5). (F) Fold changes in gene expression in SAHA- and DMSO-treated cells, showing that genes with high RNAPII binding (blue) are repressed by SAHA, as compared to other highly expressed genes (gray). Asterisks indicate p-values < 0.05.

We further observed the differential binding of transcriptionally active RNAPII, with C-terminal domain phosphorylations at Ser5 and Ser2 (S5P and S2P) in its largest subunit (Figures 2B and S2). The RNAPII binding levels decreased in the down-regulated genes, increased in the up- regulated genes, and remained the same in the unchanged genes, reflecting each transcription response to SAHA. Downregulated genes exhibited higher RNAPII binding as compared to the up- regulated and unchanged genes, under basal conditions (Figure 2B). This difference was not due to DNA amplifications, as the input reads were consistent. We then hypothesized that high RNAPII binding is a characteristic of genes susceptible to SAHA-induced repression. To test this, we identified 119 genes with highly dense RNAPII binding, by calculating the RNAPII peak scores from RNAPII (S2P) ChIP-seq data of untreated MCF-7 cells (Figure 2C). We used the S2P form, which marks elongating RNAPII, to assess occupancy across the gene body. Although these genes were highly expressed (Figure 2D), they accounted for only 6% of all highly expressed genes (Figure 2E), indicating that dense RNAPII binding is not a common feature of highly expressed genes. This suggests that dense RNAPII binding may reflect additional regulatory mechanisms beyond transcriptional output. In fact, these genes tended to be down-regulated by SAHA (Figure 2F), suggesting that highly dense RNAPII binding likely represents a chromatin status susceptible to transcriptional repression by SAHA.

### RNAPII is dissociated from the promoters of genes down-regulated by SAHA

To investigate how SAHA reduces RNAPII binding at specific genes, we conducted a genome-wide time-course analysis of RNAPII (S5P) binding and nascent RNA production by ChIP-seq and GRO-seq at 30-minute intervals. At the representative SAHA-down-regulated gene *ESR1* (300 kb), RNAPII binding was reduced at the 5′ promoter region within 30 minutes, with the reduction gradually extending toward the 3′ end over 60–90 minutes (Figure 3A). GRO-seq showed a similar decline in nascent RNA, beginning at the 5′ end and spreading downstream (Figure 3B). This progressive repression pattern was also seen in long SAHA-down-regulated genes such as *GFRA1* (216 kb) and *GREB1* (103 kb) (Figures S3A and S3B), whose lengths enabled resolution of transcriptional dynamics given typical elongation rates (>1 kb/min)^40^. Total RNA-seq across *ESR1* introns also revealed a time-dependent decline in read counts from the 5′ to 3′ direction (Figure 3C). These results suggest that SAHA primarily inhibits RNAPII recruitment to promoters, rather than disrupting RNAPII pausing or elongation (Figure 3D).

**Figure 3.**
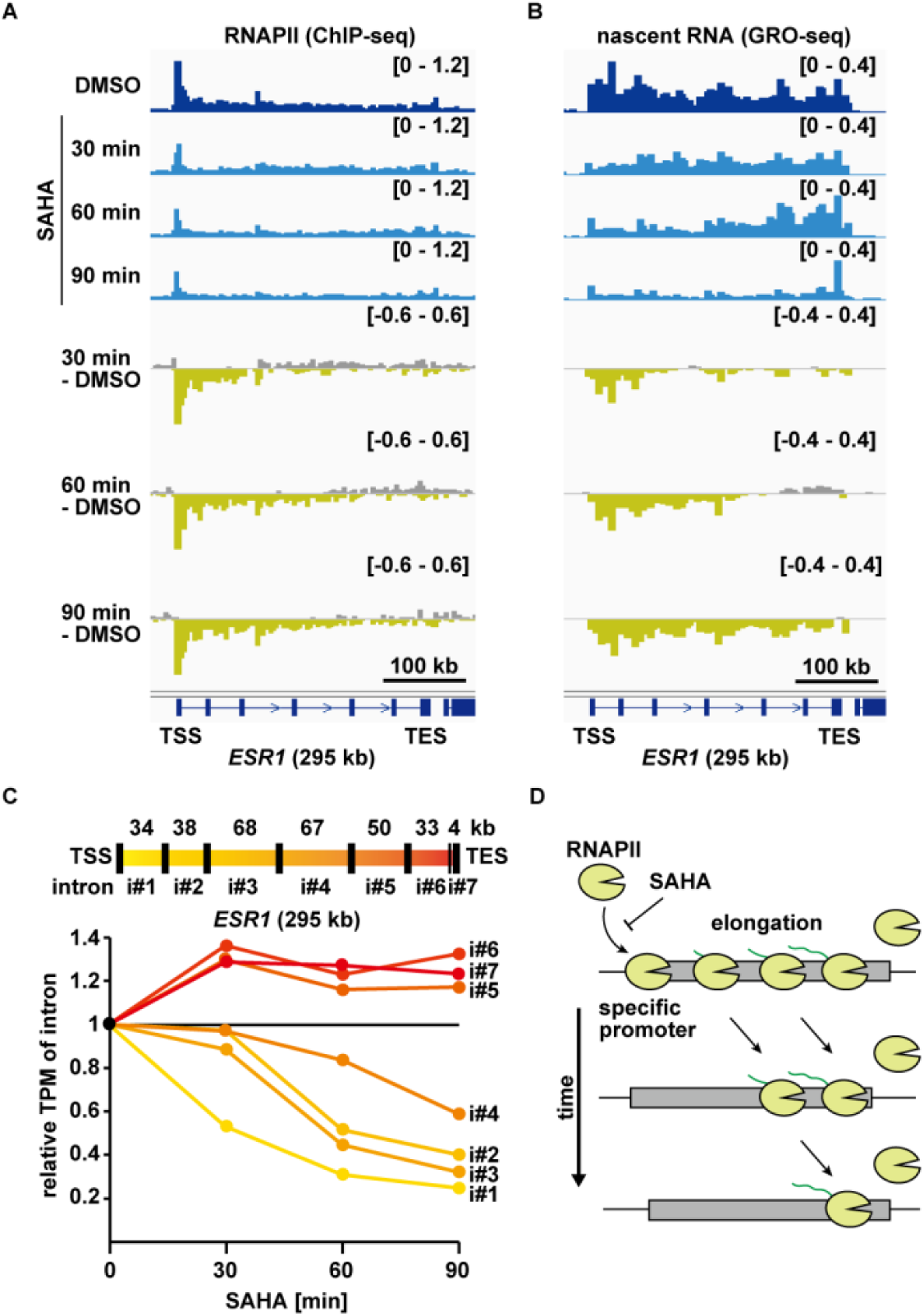
SAHA prevents RNAPII binding to the promoters of down-regulated genes, resulting in transcription inhibition. (A) Time-course analysis of RNAPII (S5P) ChIP-seq at 30-min intervals following SAHA treatment. Changes in RNAPII binding (SAHA vs. DMSO) along the ESR1 gene are visualized. Yellow areas indicate decreased binding and gray areas indicate increased binding. The reduction of RNAPII binding starts at the TSS and expands toward the TES over time. (B) Time-course analysis of GRO-seq as in (A), showing a similar reduction pattern. (C) Intronic RNA-seq signals from the *ESR1* gene over time after SAHA treatment. Values relative to those in untreated cells are shown. The extent of the transcriptional decrease was greater in introns closer to the TSS. Introns are numbered based on their proximity to the TSS. (D) A model of a SAHA-down-regulated gene. SAHA prevents RNAPII binding to the promoter, while allowing transcription elongation to continue.

### SAHA represses a distinct enhancer subset termed HD enhancers

RNAPII loading at promoters is typically mediated by multiprotein assemblies, including the transcription preinitiation complex, which also enables enhancer–promoter communication. To investigate how SAHA represses RNAPII loading, we first sought to identify enhancers within regions where RNAPII dissociates upon deacetylase inhibition. Based on changes in RNAPII occupancy following SAHA treatment compared to DMSO, we categorized 20,830 RNAPII-bound regions into five groups: increased binding (r#1 and r#2), unchanged (r#3), and decreased binding (r#4 and r#5) (Figure 4A). Genomic distribution analysis revealed that unchanged (r#3) and down- regulated regions (r#4 and r#5) were enriched in intergenic areas, which include enhancers, whereas up-regulated regions were mainly promoter-proximal (Figure 4B).

**Figure 4.**
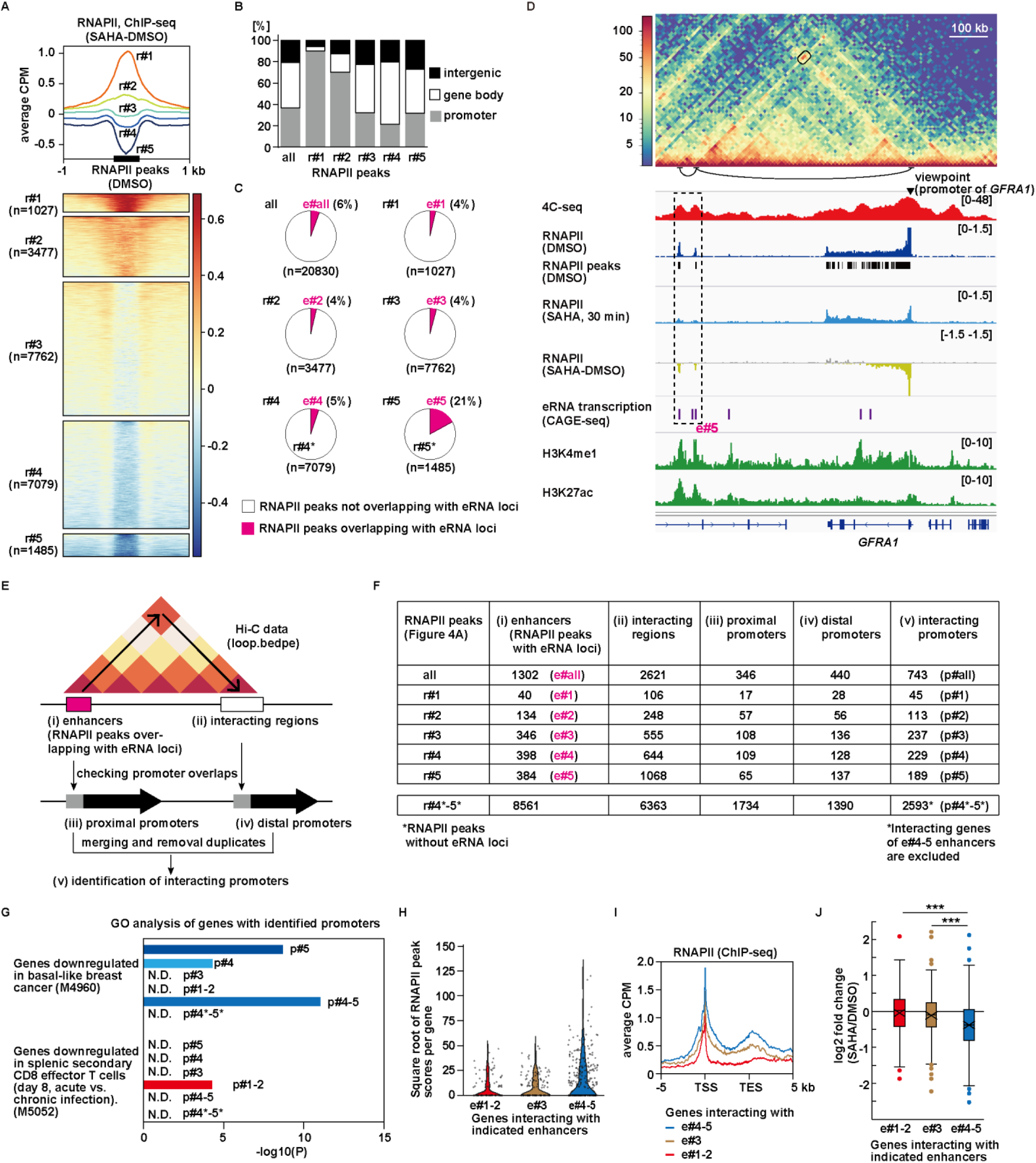
Identification of HD enhancers based on bidirectional enhancer RNA transcription and loss of RNAPII binding by SAHA. (A) Classification of RNAPII (S5P) binding regions. Differential ChIP-seq signals (SAHA minus DMSO) are shown in a plot (top) and heatmaps (bottom). The positions of RNAPII peaks in DMSO-treated MCF-7 cells are centered. Regions were divided using k-means clustering into five groups: increased (r#1, r #2), unchanged (r#3), or decreased (r#4, r#5) RNAPII binding after SAHA treatment. (B) Genomic locations of regions r#1-5 defined in Figure 4A. Promoter and gene body regions were defined based on the genomic coordinates for actively transcribed genes (TPM > 1). (C) The proportion of enhancer candidates (magenta, e#1-5) among RNAPII binding regions (r#1- 5). Enhancers were defined as sites exhibiting bidirectional eRNA transcription in the CAGE-seq analysis. We identified e#4 and e#5 as HD enhancers. (D) Representative e#5 regions near the *GFRA1* locus. Alignment of the Hi-C contact map, 4C-seq and ChIP-seq profiles revealed a special enhancer candidate region, as denoted with a dotted box. From the top, the region interacts with the *GFRA1* promoter, as shown in the Hi-C and 4C-seq profiles, and exhibits reduced RNAPII binding upon SAHA treatment (RNAPII ChIP-seq) as well as bidirectional transcription (eRNA, CAGE-seq). It is also enriched with general enhancer markers, H3K4me1 and H3K27ac. (E) Scheme for the identification of a target gene candidate for each enhancer, using Hi-C data. (F) Summary of the promoter-enhancer interactions for e#1–5. r#4* and r#5* represent regions with RNAPII dissociation upon SAHA treatment, as in e#4-5, but without eRNA transcription. (G) GO analysis using MSigDB (GSEA), showing that the genes interacting with e#4-5 are enriched with those down-regulated in basal breast cancer (top). In contrast, the genes interacting with e#1-2 are enriched with those irrelevant to luminal breast cancer, and down-regulated in splenic secondary CD8 effector T cells (bottom). (H) RNAPII peak scores (square root-transformed) for genes interacting with the enhancers e#1–5. RNAPII binding is higher in the HD enhancer (e#4-5) target genes than in others. Scores were calculated as in Figure 2C. (I) RNAPII aggregation plots reveal higher binding at the HD enhancer (e#4-5) target genes than at those of other groups, based on the ChIP-seq data from untreated MCF-7 cells used in Figure 2A. (J) Fold changes in gene expression after the SAHA treatment. The HD enhancer (e#4-5) target genes were mostly repressed by SAHA. Asterisks indicate p-values < 0.05.

To assess enhancer activity at these regions, we examined eRNA production based on bidirectional TSSs identified by CAGE-seq. The enhancers were classified into e#1–e#5, corresponding to r#1–r#5 (magenta sectors in Figure 4C). Notably, r#5, comprising regions with the highest degree of RNAPII dissociation upon SAHA treatment, showed the highest proportion of regions with eRNA transcription, suggesting that SAHA-sensitive RNAPII dissociation marks a distinct subset of active enhancers (Figure 4C).

*GFRA1*, a SAHA-downregulated gene encoding a luminal breast cancer-associated cell surface receptor (Figure 1C)^41^, is located ∼600 kb downstream of a representative e#5 enhancer. Hi- C and 4C-seq analyses revealed physical interaction between this enhancer and the *GFRA1* promoter (Figure 4D). Similarly, the promoters of other SAHA-down-regulated genes, *CCND1* and *GATA3*, interact with e#5 enhancers (Figures S4A and S4B). In contrast, the promoters of *ZFAND3* and *TRIO*, whose transcription levels remained unchanged after SAHA treatment, were not associated with e#5 enhancers (Figures S4C and S4D). These results confirm that the chromatin interactions detected in the Hi-C data could dictate a target gene for each enhancer, as previously shown.^21–24,42,43^ We therefore utilized the Hi-C data to map the genome-wide enhancer-promoter interactions (Figure 4E). This analysis identified 158 target promoters for e#1–2 (p#1–2), 237 for e#3 (p#3), and 418 for e#4–5 (p#4–5) (Figure 4F). As a control, we identified 2,593 promoters that interact with RNAPII peak regions in r#4-5 but lack eRNA transcription (r#4*-5*, white areas in Figure 4C). To isolate the genes uniquely associated with r#4*-5*, we excluded those also interacting with e#4-5.

A GO analysis revealed that the target genes for e#4-5, but not for e#1-3, are enriched in terms associated with the luminal breast cancer subtype (Figures 4G and S4E). These results indicate that a set of enhancers (e#4-5) delineating cell identity was defined based on a combination of two criteria: susceptibility of RNAPII binding under hyperacetylation (SAHA treatment) and active enhancers with bidirectional transcription of eRNAs. Because enhancer activity of e#4 and e#5 is abolished by SAHA-induced acetylation, and thus rely on hypoacetylation, we designated them as hypoacetylation-defined enhancers (HD enhancers; Table S1, genomic coordinates; Table S2, target genes).

Further analyses of the RNAPII ChIP-seq and RNA-seq datasets confirmed that the characteristics of the SAHA-down-regulated genes were recapitulated by the HD enhancer (e#4-5)- targeted genes (Figures 4H-J). The RNAPII binding levels at the gene loci and TSSs are high under basal conditions (Figures 4H and 4I). The transcription levels of the HD enhancer-targeted genes are high, but only 3% of highly expressed genes are associated with HD enhancers (Figures S4F and S4G). In addition, the transcription of HD enhancer-targeted genes is down-regulated by SAHA (Figure 4J). These findings highlight the features of a distinct subset of enhancers, HD enhancers, in regulating luminal breast cancer-associated genes.

### HD enhancers partially overlap with super-enhancers, but are distinct

Super-enhancers often contain a cluster of enhancers regulating genes critical for cell identity, like the HD enhancers we identified (Figures 4G and S4E) ^30,44–46^. To compare HD enhancers and super-enhancers, we analyzed 782 HD enhancers (e#4-5) and 390 super-enhancers identified in MCF-7 cells by H3K4me1 and H3K27ac ChIP-seq (Figure 5A). 286 of the 782 HD enhancers (36.6%) overlapped with 115 super-enhancers (common enhancers), averaging 2.5 HD enhancers per super-enhancer. For instance, three HD enhancers of *GFRA1* are located within one super-enhancer (Figure 5B, middle). Conversely, 496 HD enhancers and 275 super-enhancers did not overlap, as seen at the *TBL1XR1* and *GAN* loci, respectively (Figure 5B).

**Figure 5.**
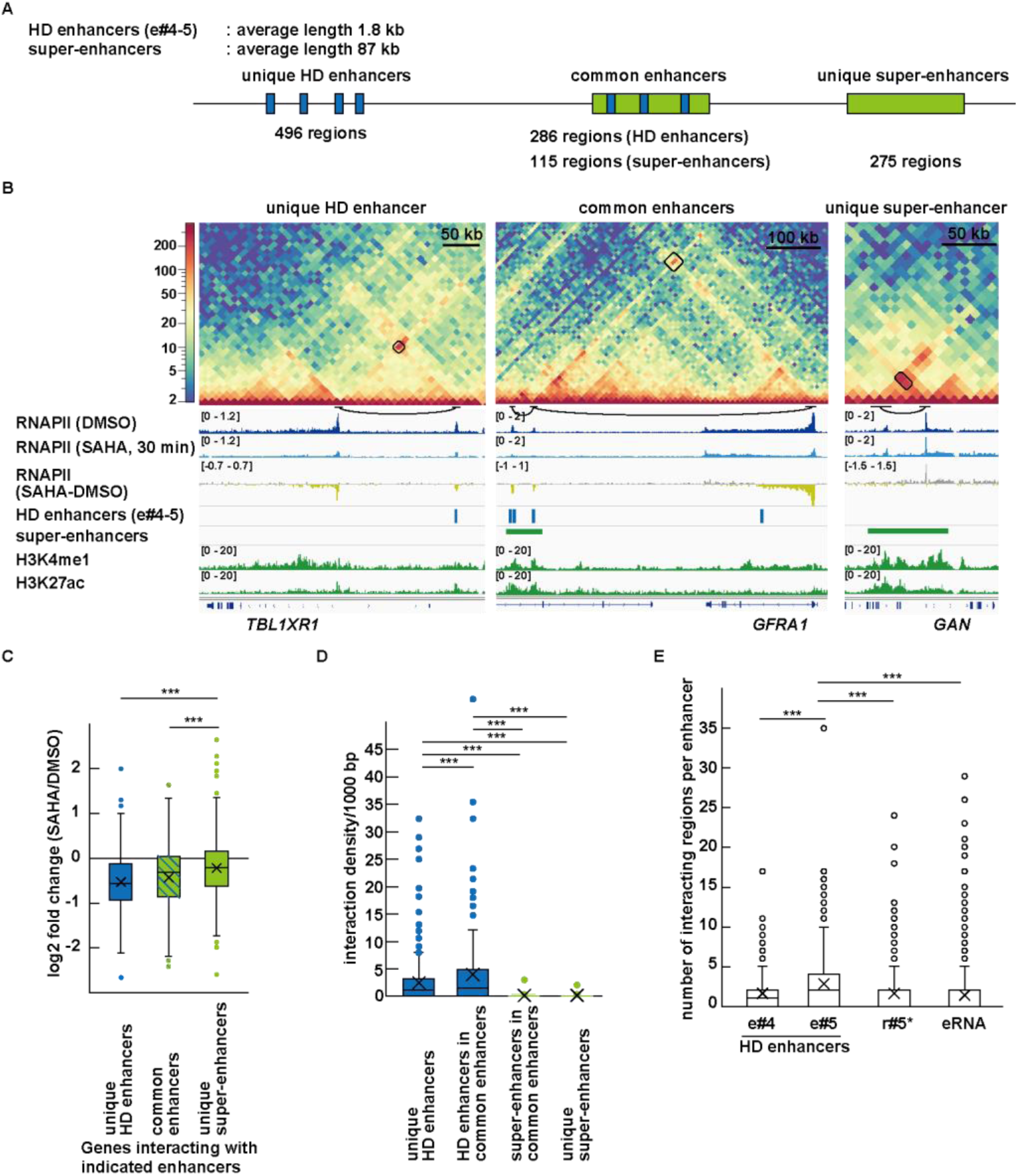
Unique and common features of HD enhancers and super-enhancers. (A) Overlap between HD enhancers (e#4-5) and super-enhancers. In MCF-7 cells, 496 unique HD enhancers (e#4, 5), 275 unique super-enhancers, and 115 common enhancers (overlapping with 286 HD enhancers (e#4-5)) were found. Due to its length, a single SE often contains multiple HD enhancers. Super-enhancers were defined based on the ChIP-seq data for H3K4me1 and H3K27ac. (B) Representative unique HD enhancers (left), common enhancers (middle), and unique super-enhancers (right). Hi-C contact maps depict promoter-enhancer interactions, and ChIP-seq data illustrate chromatin properties. (C) Fold changes in expression levels of genes interacting with the indicated enhancers, 3 h after SAHA treatment. Genes interacting with unique HD enhancers and common enhancers were down- regulated. Asterisks indicate p-values < 0.05. (D) Unique and common HD enhancers exhibit frequent chromatin interactions. The interaction density per 1,000 bp was calculated from the MCF-7 Hi-C data by dividing the number of interactions by the enhancer length. Asterisks indicate p-values < 0.05. (E) SAHA sensitivity and enhancer RNA production are associated with inter-chromatin interaction frequencies. HD enhancers with the greatest sensitivity to SAHA for RNAPII dissociation (e#5) have more chromatin-interacting regions than those with less sensitivity (e#4). Regions with similar sensitivity but no eRNA production (r#5*) have low chromatin interaction frequencies. Similarly, eRNA-producing regions, excluding e#4-5 (eRNA), do not exhibit particularly frequent interactions. Asterisks indicate p-values < 0.05.

We used Hi-C data to identify target genes for each enhancer type, as done for e#1-5 (Figure 4E). Genes interacting with HD enhancers or common enhancers were more repressed by SAHA than those interacting with unique super-enhancers (Figure 5C), indicating HD enhancer susceptibility to hyperacetylation-induced repression.

To assess the potential differences in 3D genome organization between HD enhancers and super-enhancers, we next analyzed the genome-wide chromatin interactions in MCF-7 cells, using Hi-C data. The HD enhancers showed higher interaction frequencies than the super-enhancers (Figure 5D). The HD enhancers within super-enhancers exhibited even higher interaction frequencies than the unique HD enhancer regions. Among the HD enhancers (e#4 and e#5), the regions with strong RNAPII dissociation upon SAHA treatment (e#5) exhibited greater numbers of chromatin interactions than those with weaker dissociation (e#4) (Figure 5E). Furthermore, the regions with pronounced RNAPII dissociation but lacking eRNA transcription (r#5*) showed lower interaction numbers. Similarly, the regions with bidirectional eRNA transcription independent of RNAPII dissociation also showed lower interaction numbers. These findings define HD enhancers as unique regulators with strong chromatin connectivity. Supporting this, ChIP-seq analysis revealed that MED1, a hallmark of super-enhancers, is more enriched at common enhancers than at unique super-enhancers (Figure S5), suggesting that HD enhancer–containing super-enhancers are structurally and functionally specialized.

### HD enhancers require Mediator components for their transcription regulation

The Mediator complex facilitates transcription pre-initiation complex assembly and promotes enhancer–promoter interactions via chromatin looping. Consistently, our ChIP-seq analyses revealed more MED1 binding to HD enhancers (e#4-5) in association with RNAPII, the transcription factor ERα, and H3K27ac, as compared to other enhancers (e#1-3) in MCF-7 cells under basal conditions (Figure 6A). The ChIP-seq data also revealed that the binding dynamics of MED1 and RNAPII after 30 minutes of SAHA treatment decreased at HD enhancers, remained stable at e#3, and increased at e#1-2 (Figure 6A), whereas ERα binding remained unchanged. These changes were exemplified by the representative HD target genes, *ESR1* and *CCND1* (Figures 6B and S6). 4C-seq revealed that HD enhancer–promoter interactions were preserved or even enhanced after SAHA treatment. This suggests that transcriptional repression results from Mediator dissociation rather than of enhancer–promoter contact (Figure 6B). Supporting this, knockdown of MED21 or MED24 in MCF-7 cells repressed the HD enhancer-regulated genes (*ESR1*, *CCND1*, *GFRA1*, *GATA3*), but not the non-HD enhancer-regulated genes (*ZFAND3*, *TRIO*, *RPS6KB1*), recapitulating SAHA’s effects (Figure 6C).

**Figure 6.**
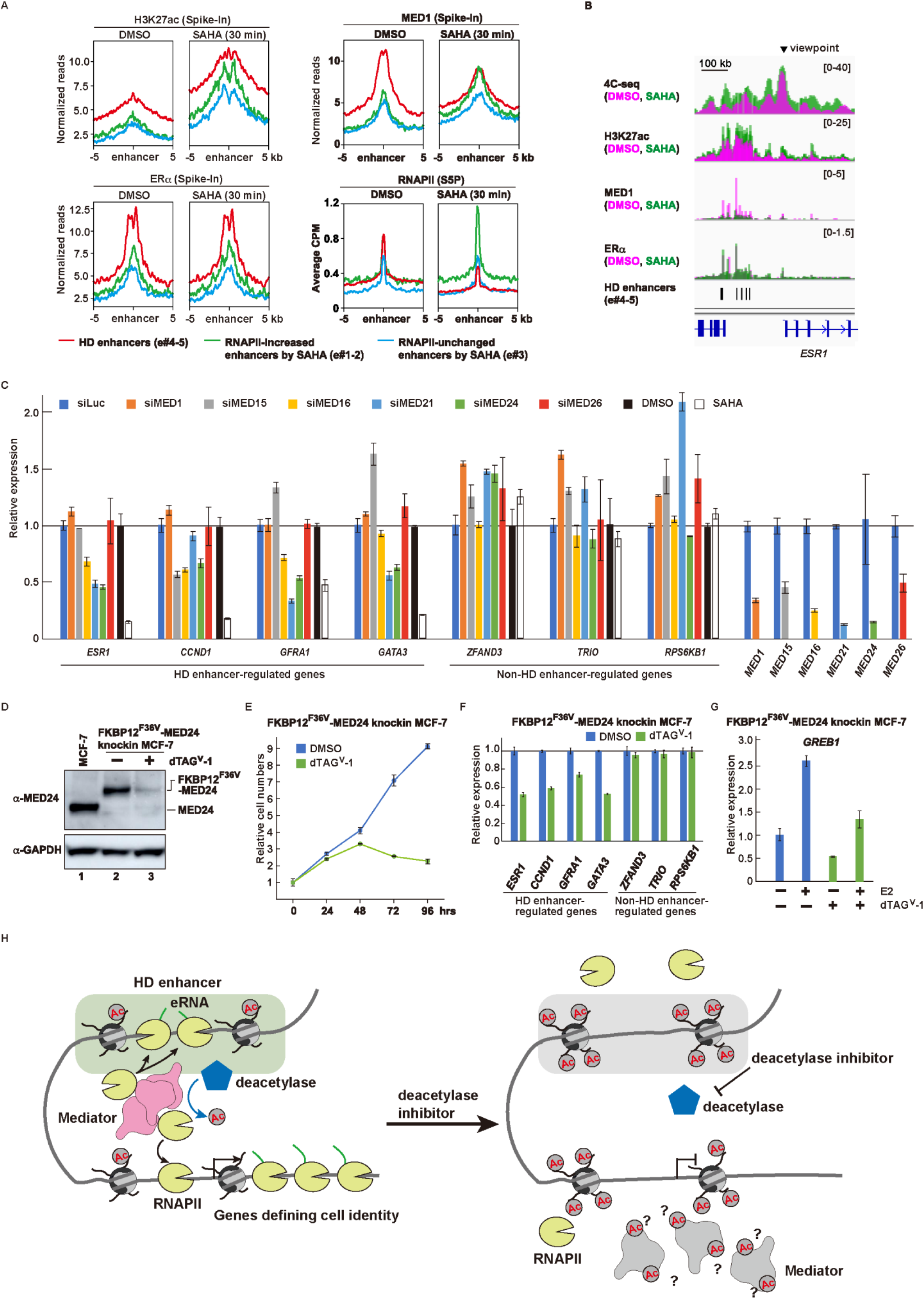
HD enhancers require the Mediator complex for their function. (A) Aggregation plots of the ChIP-seq signals around the enhancer regions. In MCF-7 cells (DMSO), the binding of H3K27ac, MED1 and Erα is higher at HD enhancers (e#4-5, red lines) than at others (e#1-2, green lines and e#3, blue lines). Upon SAHA treatment, MED1 and RNAPII were dissociated from HD enhancers, while the H3K27ac level increased across all enhancer types (red lines, DMSO vs. SAHA). Erα levels remain unchanged. (B) Representative HD enhancers near the *ESR1* gene. The 4C-seq data show that the interactions of the HD enhancers and the promoter persisted after the SAHA treatment (cyan vs. magenta). (C) Reverse transcription-PCR analysis revealed that MED21 and MED24 knockdowns specifically recapitulated the transcription repression by SAHA. Expression levels of SAHA-down-regulated genes (*ESR1*, *CCND1*, *GFRA1*, *GATA3*) and SAHA-unchanged genes (*ZFAND3*, *TRIO*, *RPS6KB1*) were analyzed following the indicated Mediator component knockdown (72 h) or SAHA treatment (6 h). (D) Western blot showing degradation of the FKBP12^F36V^-tagged endogenous MED24 protein. The MED24 protein in the parental (lane 1) and FKBP12^F36V^-MED24 knock-in (lanes 2 and 3) MCF-7 cells was detected (top). To induce specific degradation, cells were treated with the FKBP12^F36V^ degrader dTAG^V^-1 for 3 h (lane 3). GAPDH serves as a loading control (bottom). (E) Cell growth curves of FKBP12^F36V^-MED24 knock-in MCF-7 cells with (-, blue) or without (+, green) dTAG^V^-1 treatment. MED24 depletion by dTAG^V^-1 treatment repressed cell growth. (F) RT-qPCR analysis showing gene expression changes with MED24 degradation. Upon MED24 depletion by the 6 h dTAG^V^-1 treatment, HD enhancer-regulated genes (*ESR1*, *CCND1*, *GFRA1*, *GATA3*) were repressed, whereas non-HD enhancer-regulated genes (*ZFAND3*, *TRIO, RPS6KB1*) maintained their expression. (G) RT-qPCR analysis of an estradiol (E2)-induced gene, *GREB1*, in FKBP12^F36V^-MED24 knock-in MCF-7 cells. Cells were cultured in E2-free medium, treated with dTAG^V^-1 at 45 h, and stimulated with E2 at 48 h. *GREB1* expression levels were measured 3 h after E2 addition. MED24 depletion suppressed E2-induction of *GREB1* expression. (H) Model for HD enhancer regulation of cell identity genes. HD enhancers are defined as sites where eRNAs are transcribed and RNAPII is dissociated by SAHA treatment. In MCF-7 cells, HD enhancers are rich in H3K27ac and bound by the Mediator complex, which supports RNAPII recruitment to an enhancer and a promoter (black arrows). Deacetylation of non-histone protein(s) (a Mediator component, for example) by lysine deacetylase may be critical to maintain active transcription (blue arrow). Upon SAHA treatment, the deacetylation status is inhibited, and the Mediator complex and RNAPII dissociate from HD enhancers, leading to transcriptional repression.

To study MED24’s role in HD enhancer regulation, we generated an MCF-7 knock-in line with endogenous MED24 tagged with FKBP12^F36V^ for dTAG-mediated degradation (Figure 6D). Degradation of MED24 by the dTAG^V^-1 degrader reduced both cell proliferation and the expression of HD enhancer–regulated genes (Figures 6E and 6F). MED24 depletion also impaired estradiol-induced transcriptional activation of *GREB1*, a hallmark of luminal breast cancer cells (Figure 6G). Although fold induction was similar in the presence or absence of MED24, basal transcription was markedly lower in depleted cells, leading to lower absolute transcript levels after hormone stimulation. These findings suggest that MED24 supports estrogen-responsive gene activation by sustaining HD enhancer activity, and that disrupting this regulation limits the transcriptional output required for full hormonal response.

In summary, we identified 784 HD enhancers that regulate the genes associated with the luminal breast cancer identity in MCF-7 cells. These enhancers are characterized by high levels of histone acetylation, bidirectional eRNA transcription, and frequent chromatin interactions. They are also densely occupied by RNAPII and the Mediator complex, and the depletion of MED24 reduced the expression of HD enhancer-regulated genes, indicating that the complex is essential for their transcriptional activity. HD enhancers are suppressed upon deacetylase inhibition, suggesting that the hypoacetylated state of their complexes is critical for their functions (Figure 6H).

### Patient Tumor Analysis Identifies HDC1, an HD Enhancer Subgroup Linked to Transcriptomic Subtypes

Using ATAC-seq data from breast cancer samples^47^, we further examined whether HD enhancer regions identified in MCF-7 cells are accessible in patient tumors, suggesting their potential enhancer activity (Figure 7A). A subset of ATAC peaks within HD enhancers (e#4-5) exhibited subtype-specific variations in signal strength, as indicated by the vertical lines on the left side of the heatmap (Figure 7B). In contrast, ATAC-seq peaks in e#1 to e#3 showed relatively consistent levels across all subtypes. A UMAP analysis using the ATAC signal intensities at each enhancer revealed that the HD enhancers separated patients by subtypes, most notably basal tumors clustering from other subtypes (Figure 7C). In line with this observation, the variance in ATAC-seq signals was significantly higher in the HD enhancers (e#4-5) than in the other enhancer groups, suggesting that a distinct subset of HD enhancers is responsible for subtype-specific gene expression (Figure 7D).

**Figure 7.**
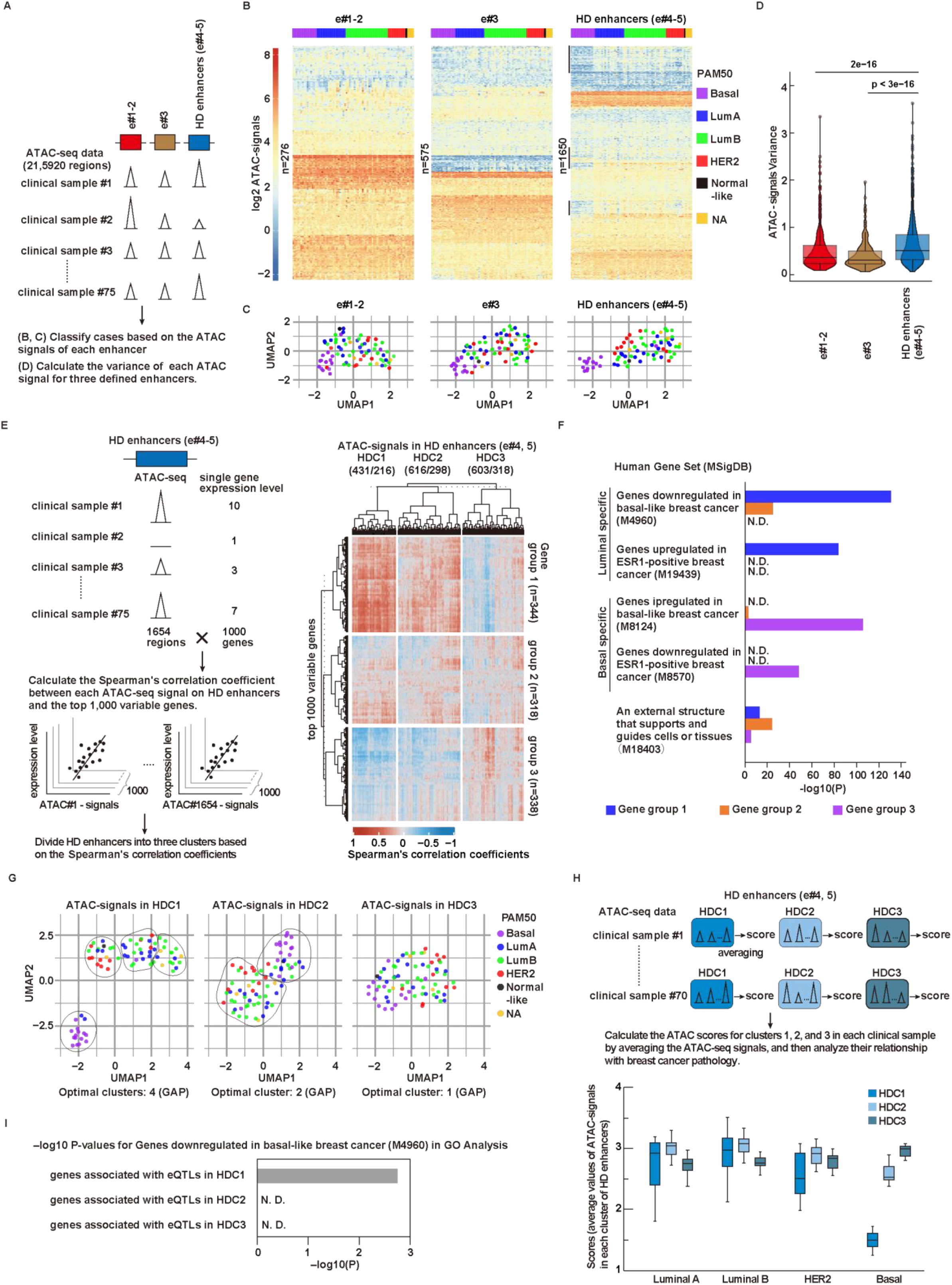
HDC1 enhancers, a subpopulation of HD enhancers, reflect transcriptome-based clinical subtypes. (A) Scheme to map each enhancer in e#1-5 to an ATAC-seq signal in primary breast tumors (n=75) from the TCGA database. For each clinical case, transcriptome (PAM50)-based subtypes are annotated as basal (purple, n=15), HER2 (red, n=11), luminal A (blue, n=18), luminal B (green, n=26), Normal-like (black, n=1), and Not Available (NA, gray, n=4). (B) The alignment of ATAC signals according to PAM50-defined subtypes revealed a subset of ATAC-peaks in HD enhancers (e#4-5) showing differential signal strengths (black lines). In contrast, most ATAC signals corresponding to other enhancers (e#1-3) were relatively constant, regardless of subtype differences. Regions within HD enhancers that exhibited marked signal variation among subtypes are indicated by black lines alongside the heatmap. (C) UMAP analysis revealing that HD enhancers (e#4-5) can discriminate basal cases from others effectively. (D) Violin plot showing the variance of ATAC signals corresponding to enhancers (e#1-5). HD enhancers (e#4-5) exhibit higher variation. (E) A strategy to classify the HD enhancers (e#4-5) based on the correlations between their ATAC- seq signal strength and gene expression levels in clinical samples (left). The top 1,000 genes with the largest transcriptional variability across cases were selected for analysis. HD enhancers (e#4-5) were classified into three groups, HDC1 (HD Cluster1), HDC2, and HDC3 (right). The 1,000 genes were also grouped into Gene groups 1, 2, and 3, based on the dendrogram on the left of the panels (vertical dotted line). (F) GO analysis using MsigDB (GSEA), showing that Gene group 1 (blue bars) is enriched in luminal-specific gene sets, and gene group 3 (purple bars) is enriched in basal-specific gene sets. (G) The UMAP analysis of ATAC-seq signals at HDC1 enhancer regions divides cases into four groups that largely correspond to PAM50 subtypes (basal, HER2, luminal A, luminal B), whereas HDC2 and HDC3 yielded more homogeneous distributions. Cluster numbers were determined by Gap statistics. (H) Strategy to calculate average intensities for ATAC signals corresponding to HDC1-3 (top). The signals for HDC1 are high in the luminal A, luminal B, and HER2 subtypes, but low in the basal type patients (bottom). Additionally, HDC1 shows greater variation within the same subtype (luminal A, luminal B, HER2), as compared to clusters 2 and 3. Cases classified as Normal-like or Not Available were omitted in this analysis, due to their small sample sizes (n=1 and 4, respectively). (I) The GO enrichment analysis indicates that the eQTLs in HDC1 enhancers are associated with luminal genes, whereas the eQTLs of HDC2 and HDC3 are not. The bar plots display the top enriched biological processes for each gene set. Enrichment scores are shown as –log_10_ adjusted p-values.

To investigate how HD enhancers are involved in subtype-specific gene expression, we analyzed the correlations between the ATAC signal intensities within HD enhancers and the expression levels of the top 1,000 genes with variable expression across patient tumors (Figure 7E, left). With the correlation pattern based on Spearman’s correlation coefficients, the HD enhancers were divided into three clusters, which we termed the HDC1 (HD Cluster 1), HDC2 and HDC3 enhancers (Figure 7E, right; Table S3, correlation matrix; Table S4, genomic coordinates of each cluster). The HDC1 and HDC2 enhancers are positively correlated with Gene group 1, moderately correlated with Gene group 2, and negatively correlated with Gene group 3, with HDC2 displaying a less pronounced correlation. HDC3 exhibited a reversed and less obvious correlation. According to GO analysis, HDC1 enhancer formation correlates positively with luminal-associated genes and negatively with basal-associated genes, whereas HDC2 and HDC3 show weak or no subtype-specific correlations (Figure 7F). To confirm the HDC1 enhancers as subtype-defining enhancers, we identified the candidate target genes for HDC1 using the Hi-C data of MCF-7 cells, and compared them with the other gene sets analyzed in this study (Table S5). Comparisons of the P-values on the GO term of luminal specific gene set showed that the genes interacting with HDC1 enhancers are most enriched in this category (Figure S7).

To assess whether breast cancer cases could be stratified with HD enhancer clusters, we performed a UMAP analysis on the ATAC-seq signals from HDC1-3 enhancers and calculated the GAP statistic (Figure 7G). While HDC2 and HDC3 failed to classify cases by clinically relevant subtypes, HDC1 stratified patients into four distinct groups. One group, predominantly composed of basal cases, also included two luminal A cases. A cluster associated with the HER2 subtype contained both luminal A and B cases. The groups corresponding to luminal A and B were partially intermingled. These analyses revealed that HDC1 chromatin accessibility only partially aligns with PAM50 subtypes. These discrepancies may reflect the inherent heterogeneity and plasticity of breast cancer biology, which cannot be fully captured by transcriptome-based classifications alone. Alternatively, the HDC1-based chromatin features may reveal previously unrecognized regulatory subtypes, highlighting the potential of enhancer-defined markers as complementary tools for patient stratification. Another interesting observation was that HDC1 displayed a broader range of ATAC signal intensity in luminal and HER2 patients (Figure 7H). This suggests variability in chromatin accessibility and in the assembly of enhancer-associated regulatory complexes at HDC1 enhancers. Such variability may underlie inter-patient differences in transcriptomic profiles and disease phenotypes, as previously described^48,49^. We further examined eQTLs derived from the GTEx Portal (GTEx Analysis V7, Breast Mammary Tissue). We found that the SNPs located in the HDC1 locus were associated with the expression of genes specifically expressed in luminal breast cancer (Figure 7I). This result supports the notion that HDC1 may function as a subtype-specific regulatory element.

In summary, integrative chromatin analyses of breast cancer cell lines and patient tumors identified a distinct enhancer subset, HDC1, that may play a critical role in defining breast cancer subtype identity.

## Discussion

In this study, we identified 784 HD enhancers that are associated with genes that define luminal breast cancer cell identity. Importantly, the HD enhancers of the *ESR1* and *CCND1* genes were colocalized with transcriptional regulatory elements detected by genome-scale and single-cell CRISPR screens^39^, underscoring their functional significance. By analyzing clinical datasets, we further refined this set to 216 HDC1 enhancers whose chromatin accessibility stratifies transcriptome-defined breast cancer subtypes. This specific set of enhancers may play a crucial role in shaping global transcriptional patterns, reflecting both the etiology and therapeutic responsiveness of breast cancer.

A notable feature of genes associated with HD enhancers is their dense RNAPII occupancy, both at transcription start sites and along gene bodies. While these genes are highly expressed, this pattern of dense RNAPII binding is not observed in all highly expressed genes. This suggests that dense RNAPII occupancy may reflect a distinct transcriptional mode characteristic of cell identity genes. Similar patterns have been reported in other contexts. In differentiated mouse cells, long and highly expressed genes extend across the nucleus and are decorated with progressively growing nascent ribonucleoproteins, forming so-called transcription loops^50^. An exogenously introduced HIV genome also showed dense and continuous RNAPII convoy^51^. Moreover, elevated RNAPII accumulation has been detected at histone gene clusters undergoing hypertranscription in cancer patient samples, correlating with centromeric breaks and whole-arm chromosomal imbalances-events known to drive tumorigenesis, and proposed as a prognostic marker in human cancers^52^.

In addition to their dense RNAPII occupancy, genes associated with HD enhancers exhibit marked RNAPII loss upon lysine deacetylase inhibition, highlighting their unique transcriptional regulation. HDACs are widely mapped to transcriptionally active genomic loci. It has been proposed that histone deacetylation promotes RNAPII engagement in transcriptional cycles, including initiation, termination, and re-initiation, through repeated cycles of histone acetylation and deacetylation^31,53^. A recent study demonstrated that p300/CBP-mediated acetylation of Mediator components such as MED1 is enhanced by HDAC inhibition^54^. This acetylation impairs transcriptional condensate formation and RNAPII interactions via the CTD, potentially repressing transcription (Figure 6D). Our analysis suggests that MED21 and MED24, but not other Mediator components, are required to maintain cell identity gene expression. This may reflect distinct Mediator complex compositions at HD enhancers, consistent with prior findings implicating MED21 and MED24 in transcriptional regulation and breast tissue specificity, respectively^55,56^.

The HDC1 enhancers we identified in this study well reflect the PAM50-defined transcriptomes and breast cancer subtypes. The eQTL analysis revealed that these enhancers are significantly associated with luminal subtype-specific gene expression. UMAP analysis revealed that HDC1 clearly separates most basal tumors, while luminal subtypes showed greater heterogeneity, with some cases clustering alongside the HER2 group (Figure 7G). These observations suggest that the HDC1 enhancers capture the heterogeneity of enhancer activity and the diverse regulatory programs in luminal subtypes. In line with this, a large-scale analysis of over 1,000 luminal A tumors revealed that this subtype can be further divided into four molecular subtypes, including the "atypical luminal A," which is characterized by TP53 mutations, Aurora kinase activation, and poor prognosis^49^. Additionally, PAM50-based transcriptomic analyses have proposed a further classification into "HER2-amplified luminal" and "conventional luminal" subtypes, with the former exhibiting markedly worse clinical outcomes^48^. These findings highlight the transcriptional and genomic heterogeneity within luminal A tumors, which has direct clinical implications. Our results suggest that HDC1 enhancers may serve as epigenetic markers to capture this diversity.

While this study focuses on the luminal breast cancer subtype, the mechanisms involving HD enhancers may be shared with the enhancer sets for other breast cancer subtypes and even other diseases and biological aspects. By applying the analytical framework established here to identify HD enhancers, it may be possible to discover context-specific enhancer sets in various cell types and cancer models. Our study lays the groundwork for cell identity-defining enhancers, paving the way for deeper understanding of cancer biology and fundamental cellular processes.

## Methods

### Cell culture

The luminal breast cancer cell line MCF-7 was cultured in RPMI 1640 medium, supplemented with 10% fetal bovine serum and 1% Penicillin-Streptomycin-Glutamine mixed solution (Nacalai, 06168-34), at 37°C in a 5% CO_2_ atmosphere.

### Western blotting

MCF-7 cells were treated with 2 μM SAHA for the time periods indicated in the figure legend, washed with PBS, and detached using a cell scraper. The cells were collected by centrifugation at 3,000 × g for 5 min, and whole-cell extracts were prepared by lysing them in 1× SDS sample buffer (63 mM Tris-HCl, pH 6.8, 10% glycerol, 2% SDS, 50 mM DTT, 0.01% bromophenol blue).

Proteins were separated on a 12% NuPAGE Bis-Tris gel (Invitrogen, #12590137) using MES running buffer and subsequently transferred onto a 0.45 μm PVDF membrane (Cytiva, # 10600029). Membranes were blocked with Blocking One-P (Nacalai, #05999-84) and incubated overnight at 4°C with the following primary antibodies: pan-H3 (MAB Institute Inc. #MABI0301, 1/1,000), H3K27ac (MAB Institute Inc. #MABI0309, 1/500), H4Kac (Merck, #06-866, 1/1,000), and H4K16ac (Active Motif #39068, 1/1,000). After washing the membranes three times with TBST (Tris-buffered saline with 0.1% Tween-20) for 10 min each, they were incubated with HRP- conjugated secondary antibodies (CST #7074 for anti-rabbit, CST #7076 for anti-mouse) for 1 h at room temperature. The membranes were then washed three times with TBST for 10 min each before detection. Detection was performed using an enhanced chemiluminescence (ECL) system (Chemi-Lumi One Super, Nacalai, #02230), and signals were captured using an Amersham Imager 680 (Cytiva).

### Chromatin immunoprecipitation and ChIP-seq

Approximately 2 × 10⁷ MCF-7 cells, either untreated or treated with 2 μM SAHA for the time periods indicated in the figure legend, were cross-linked with 1% formaldehyde in PBS for 15 min at room temperature, and the reaction was quenched by adding glycine to a final concentration of 125 mM. After an incubation for 3 min at room temperature, the cells were rinsed twice with ice-cold PBS and collected using a cell scraper. After centrifugation at 500 × g for 5 min, the supernatant was removed, and for RNAPII ChIP, the pelleted cells were lysed in 1 ml of PBS containing 0.5% IGEPAL CA-630, a protease inhibitor cocktail (Roche, #4693132001), and a phosphatase inhibitor (Nacalai, #07575-51). The lysate was then centrifuged at 500 × g for 5 min, and the pellet was resuspended in 500 μl of ChIP buffer (10 mM Tris-HCl, pH 8.0, 200 mM KCl, 1 mM CaCl₂, 0.5% IGEPAL CA-630). This step was omitted for histone ChIP, where the cells were directly resuspended in ChIP buffer after the initial centrifugation. For RNAPII, MED1, and ERα ChIP, chromatin was digested using a Bioruptor (Diagnode, #BR2006A, 30 sec ON, 30 sec OFF, Level High, 5 cycles × 3). For histone ChIP, chromatin was briefly disrupted using a sonicator (TOMY, #UD-211, output 10, 40% pulse, 20 times), followed by digestion with 30 U MNase (Worthington) at 37°C for 30 min. The reaction was then terminated by adding EDTA to a final concentration of 10 mM. For RNAPII, MED1, ERα ChIP, and histone ChIPs, the solubilized chromatin fragments were separated from the pellets by centrifugation at 16,000 × g for 5 min at 4°C. To determine the DNA concentration as a proxy for the chromatin concentration, 50 μl of the solubilized chromatin was mixed with 45 μl of ChIP elution buffer (50 mM Tris-HCl, pH 8.0, 10 mM EDTA, 25 mM NaCl, and 1% SDS) containing 5 μl of 5 M NaCl, and incubated overnight at 65°C to reverse the cross-links, while the remaining solubilized chromatin was kept on ice. The DNA from the reverse cross-linked chromatin was then eluted with 0.4 mg/ml Proteinase K and further purified using a PCR clean-up kit (MACHEREY-NAGEL). The resulting DNA concentrations were measured using a spectrophotometer. For spike-in normalization, which was performed as previously reported,^57^ one two-hundredth of Drosophila spike-in chromatin (Active Motif) was mixed with the remaining chromatin. For ChIP antibody-bead preparation, 20 μl of Dynabeads M-280 anti-mouse IgG or anti-rabbit IgG (Invitrogen) per sample was washed once with 1 ml of wash buffer (1 mg/ml BSA, 2 mM EDTA in PBS). The beads were then incubated with 2 μg of the respective antibody (with an additional 2 μg of spike-in antibody, if spike-in normalization was performed) at room temperature for 60 min with rotation. After the incubation, the beads were washed twice with 1 ml of wash buffer and subsequently mixed with the chromatin samples. After gentle mixing by rotation at 4°C overnight, the beads were washed three times with 500 μl of ChIP buffer, ChIP buffer containing 500 mM KCl, and TE, respectively. The beads were then resuspended in 100 μl of ChIP elution buffer (50 mM Tris-HCl, pH 8.0, 10 mM EDTA, 25 mM NaCl, and 1% SDS) and incubated overnight at 65°C to reverse the cross-links. The DNA was then eluted with 0.4 mg/ml Proteinase K and further purified using a PCR clean-up kit (MACHEREY-NAGEL). For RNAPII, MED1, and ERα ChIP, the eluted DNA was further purified using SPRIselect beads (Beckman Coulter, #B23318). The DNA was adjusted to a final volume of 50 μl by adding dH₂O, followed by the addition of 27.5 μl (0.55×) SPRIselect beads. The mixture was pipetted 10 times to ensure thorough mixing. Using a magnetic stand, 75 μl of the supernatant was collected, and an additional 93.75 μl [(1.8 - 0.55)×] of SPRIselect beads was added. The mixture was pipetted 10 times and incubated on a magnetic stand. The supernatant was discarded, and the beads were washed with 180 μl of 80% ethanol while remaining on the magnetic stand. After brief air drying, the beads were resuspended in 20 μl of TE buffer and mixed by pipetting 10 times.

Finally, using a magnetic stand, 18 μl of the supernatant was collected as the purified DNA sample. DNA libraries were prepared using a SMARTer ThruPLEX DNA-seq kit (Takara Bio). The samples were sequenced on an Illumina HiSeq X or NovaSeq X Plus system by Macrogen.

The following antibodies were used for ChIP: anti-H3K27ac (MAB Institute Inc., #MABI0309), anti-H3K4me1 (MAB Institute Inc., #MABI0302), anti-Phospho RNAPII CTD (Ser5) (MAB Institute Inc., #MABI0603), anti-Phospho RNAPII CTD (Ser2) (MAB Institute Inc., #MABI0602), anti-ERα (Sigma-Aldrich, #06-935), anti-MED1 (Bethyl Laboratories, #A300- 793A), and anti-H2Av spike-in (Active Motif, #61686).

### RNA-seq

MCF-7 cells were cultured in 6-well plates and treated with or without 2 μM SAHA for the time periods indicated in the figure legend. Total RNA was extracted using a Maxwell RSC SimplyRNA Cells Kit (Promega) and quantified using a spectrophotometer. The ribosomal RNA (rRNA) was then depleted using a NEBNext rRNA Depletion Kit V2 (Human/Mouse/Rat) (New England Biolabs). Following rRNA depletion, RNA-seq libraries were prepared using a NEBNext Ultra II Directional RNA Library Prep Kit for Illumina (New England Biolabs), according to the manufacturer’s instructions. The samples were sequenced on an Illumina NextSeq 550 system.

### GRO-seq

GRO-seq was performed as previously reported with modifications.^58^ MCF-7 cells were grown in two 15 cm dishes per condition and treated with 2 μM SAHA for 0, 30, 60, and 90 min. After the incubation, the cells were rinsed with PBS, treated with trypsin, and collected by adding 8 ml of RPMI containing 2 μM SAHA or DMSO (for 0 min control). The cells were pelleted by centrifugation at 600 × g for 5 min, the supernatant was discarded, and the pellet was rinsed with 10 ml of ice-cold PBS. The cell pellet was resuspended in 10 ml of swelling buffer (10 mM Tris-HCl, pH 7.5, 2 mM MgCl₂, 3 mM CaCl₂, 2 U/ml Superase-In (Invitrogen)) and incubated on ice for 5 min. The cells were pelleted by centrifugation at 600 × g for 5 min, and resuspended in 10 ml of swelling buffer. This washing step was repeated once more. The pellet was then resuspended in 10 ml of lysis buffer (10 mM Tris-HCl, pH 7.5, 2 mM MgCl₂, 3 mM CaCl₂, 10% glycerol, 1% IGEPAL, 2 U/ml Superase-In) and incubated on ice for 5 min. The nuclei were pelleted by centrifugation at 600 × g for 5 min and washed twice with 10 ml of lysis buffer. After the final centrifugation, the pellet was resuspended in 3 ml of freezing buffer (40% glycerol, 5 mM MgCl₂, 0.1 mM EDTA, 50 mM Tris-HCl, pH 8.3), and the nuclei were counted. The nuclei were pelleted by centrifugation at 600 × g for 5 min and resuspended in 100 μl of freezing buffer to achieve a final concentration of 2 × 10⁸ nuclei/ml. The nuclei were stored at -80°C until use. For the run-on reaction, 100 μl of thawed nuclei was mixed with an equal volume of pre-warmed run-on reaction buffer (10 mM Tris-HCl, pH 8.0, 5 mM MgCl₂, 300 mM KCl, 1 mM DTT, 500 μM ATP, 500 μM GTP, 500 μM 4-thio-UTP, 2 mM CTP, 200 U/ml Superase-In, and 1% Sarkosyl (N-laurylsarcosine sodium salt solution)). The mixture was pipetted 15 times with a cut-tip to ensure thorough mixing and incubated at 30°C for 7 min.

To extract RNA, 600 μl of TRIzol reagent was added, and the mixture was vortexed thoroughly and incubated at room temperature for 5 min. Next, 160 μl of chloroform was added, mixed vigorously by hand for 15 sec, and incubated at room temperature for 2 min. The mixture was centrifuged at 20,000 × g for 15 min at 4°C, and the upper aqueous phase was transferred to a new tube. To precipitate RNA, 400 μl of isopropanol was added, and the mixture was incubated at room temperature for 10 min, followed by centrifugation at 20,000 × g for 10 min at 4°C. The RNA pellet was washed with 1 ml of 75% ethanol, centrifuged at 20,000 × g for 10 min at 4°C, air-dried, and resuspended in 100 μl of nuclease-free water. The RNA concentration was measured. A total of 150 μg of RNA in 500 μl of nuclease-free water was fragmented using a Bioruptor UCD-200 (Diagenode) for 1 cycle of 30 sec ON, 30 sec OFF at high power settings. Fragmentation efficiency was analyzed using a Bioanalyzer RNA Pico Kit (Agilent Technologies), following the manufacturer’s instructions. If needed, the fragmented RNA can be stored at -80°C. The fragmented RNA was then incubated at 65°C for 15 min, and cooled on ice for 5 min. Next, 500 μl of Biotinylation Solution (20 mM Tris-HCl, pH 7.5, 2 mM EDTA, pH 8.0, 40% dimethylformamide, 200 mg/ml EZ-Link HPDP Biotin (Thermo Scientific)) was added, and the RNA was biotinylated for 2 h at 25°C in the dark with shaking at 800 rpm. To purify RNA, including biotin-labeled RNA, 800 μl of chloroform was added, and the mixture was centrifuged at 20,000 × g for 5 min at 4°C. The upper aqueous phase was transferred to a new tube, and 100 μl of 5 M NaCl and 1 ml of isopropanol were added and mixed. The RNA was precipitated by centrifugation at 20,000 × g for 5 min at 4°C, and after the pellet was washed with 1 ml of 75% ethanol, it was centrifuged at 20,000 × g for 30 min at 4°C. The pellet was resuspended in 200 μl of nuclease-free water. The resuspended RNA was then incubated at 65°C for 15 min, and cooled on ice for 5 min. To isolate the biotinylated RNA, 100 μl of Dynabeads M-280 Streptavidin (Invitrogen) was pre-washed twice with 200 μl of Wash Buffer (100 mM Tris-HCl, pH 7.5, 10 mM EDTA, pH 8.0, 1 M NaCl, 0.1% (v/v) Tween-20), resuspended in 100 μl of Wash Buffer, and added to the RNA solution. The mixture was incubated at 4°C for 15 min with rotation. The beads were placed on a magnetic stand and washed three times with 900 μl of pre-warmed Wash Buffer at 65°C. The beads were then washed three additional times with 900 μl of Wash Buffer at room temperature. To elute the biotin-labeled RNA, the beads were resuspended in 100 mM DTT and incubated at room temperature for 5 min. The beads were then placed on a magnetic stand, and the supernatant was collected. This DTT elution step was repeated once more, and a total of 200 μl of eluate was collected. The eluted RNA was purified using an RNA Clean & Concentrator Kit (ZYMO RESEARCH, #R1015) and eluted in 7 μl of nuclease-free water. The RNA concentration was measured using a QuantiFluor RNA System (Promega, #E3310).

The GRO-seq libraries were prepared using a NEBNext Ultra II Directional RNA Library Prep Kit for Illumina (New England Biolabs), according to the manufacturer’s instructions. The samples were sequenced on an Illumina HiSeq X system by Macrogen.

### 4C-seq

MCF-7 cells were cultured in three 10 cm dishes per condition and treated with DMSO or 2 μM SAHA at 37°C with 5% CO₂ for 30 min. The cells were detached using trypsin and collected in RPMI 1640 medium containing either DMSO or 2 μM SAHA. PBS was added to adjust the cell concentration to 2 × 10⁶ cells/ml. An equal volume of 4% Fixation Buffer (4% formaldehyde, 7.5% fetal bovine serum in PBS) was added to the cell suspension, and the mixture was rotated at room temperature for 10 min. To quench the fixation, 2 M glycine was added to a final concentration of 130 mM, and the sample was incubated at room temperature for 2 min. The cells were then collected by centrifugation at 500 × g for 5 min.

The cell pellet was resuspended in 1 mL of ice-cold cell lysis buffer (50 mM Tris-HCl, pH 7.5, 0.5% NP-40, 1% Triton X-100, 150 mM NaCl, 5 mM EDTA, and a protease inhibitor cocktail) and incubated on ice for 20 min. The sample was then centrifuged at 500 × g for 5 min at 4°C, and the pellet was resuspended in 500 μl of 1.2× RE1 buffer (NEB, for DpnII) and pre- warmed at 37°C. After 15 μl of 10% SDS was added, the sample was incubated at 37°C with shaking for 1 h. After this incubation, 75 μl of 20% Triton X-100 and 100 U of DpnII (NEB) were added, followed by an incubation at 37°C with shaking for 3 h. An additional 100 U of DpnII was then added, and the sample was incubated overnight at 37°C with shaking.

To confirm successful digestion, 5 μl of the sample was analyzed by agarose gel electrophoresis. After verification, the remaining sample was incubated at 65°C for 20 min to inactivate the DpnII. The sample was then transferred to a 50 ml tube, and 700 μl of 10× ligation buffer (NEB, B0202) and 5.72 ml of dH₂O were added. After 50 U of T4 DNA ligase (Roche) was added, the reaction was incubated at 16°C overnight.

To confirm successful ligation, 40 μl of the sample was analyzed by agarose gel electrophoresis. After verification, 30 μg of Proteinase K (Promega) was added to the sample, and the mixture was incubated at 65°C overnight.

The DNA was phenol-chloroform extracted and recovered by ethanol precipitation. The pellet was resuspended in 1 ml of 10 mM Tris-HCl, pH 7.5. After 900 μl of AMPure XP beads was added, the sample was incubated at room temperature for 3 min. The beads were transferred to a magnetic stand, and the supernatant was removed. The beads were washed twice with 1 ml of 70% ethanol and air-dried, and the DNA was eluted in 150 μl of 10 mM Tris-HCl, pH 7.5. The eluted DNA was then divided into two portions, and 60 U of HindIII (for *ESR1*) or 50 U of Csp6I (for *GATA3*, *CCND1*, *ZFAND3*, *TRIO*, and *GFRA1*) was added to each portion. The reactions were incubated at 37°C overnight.

After confirming successful digestion by agarose gel electrophoresis, the HindIII-treated sample was incubated at 70°C for 15 min, and the Csp6I-treated sample was incubated at 65°C for 20 min to inactivate the respective enzymes. The samples were then ligated using T4 DNA ligase (50 U) at a final DNA concentration of 5 ng/μl and incubated at 16°C overnight. The ligated DNA was precipitated with ethanol and resuspended in 500 μL of 10 mM Tris-HCl (pH 7.5). The sample was then purified using 900 μl of AMPure XP beads, as described above, and eluted in 150 μl of 10 mM Tris-HCl (pH 7.5). The DNA concentration was then measured. PCR amplification was then performed using 100 ng of DNA as a template, Expand Long Template Polymerase Mix, and gene- specific primers designed using Primer Designer for 4C Viewpoints (https://mnlab.uchicago.edu/4Cpd/). Afterwards, 40 μl of AMPure XP beads was added to purify the amplified DNA using the previously described method, and the DNA was eluted in 50 μl of 10 mM Tris-HCl (pH 7.5). A 5 μl portion of the eluted DNA was then used as a template for indexing PCR with Expand Long Template Polymerase Mix. The indexed DNA was purified using a PCR clean-up kit, and sequencing was performed on the Illumina Novaseq6000 platform.

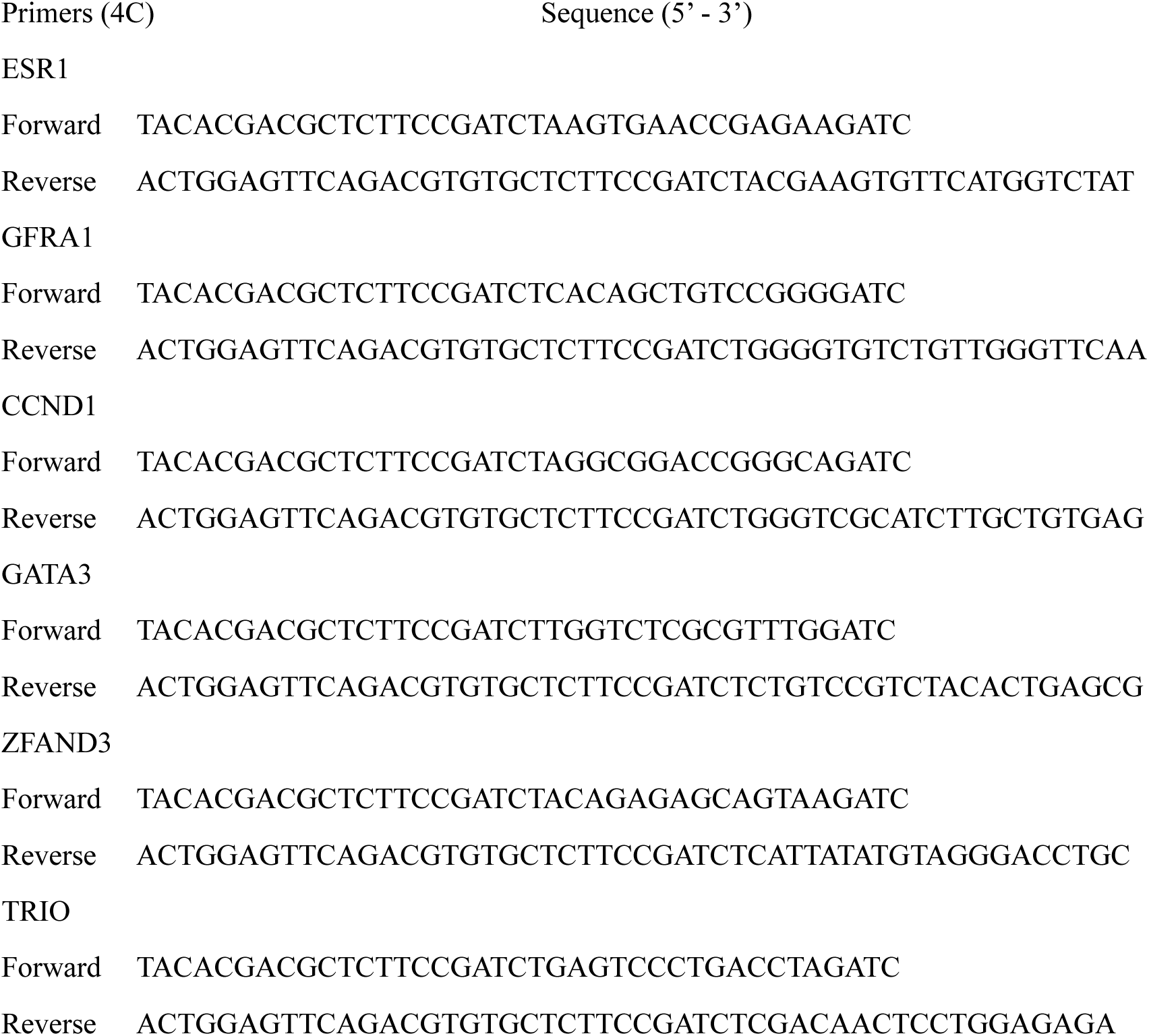

### siRNA transfections and qPCR

All siRNAs were transfected into MCF-7 cells using Lipofectamine RNAiMAX (Invitrogen), according to the manufacturer’s protocol. After 72 h, RNA was extracted using a Maxwell RSC SimplyRNA Cells Kit (Promega). cDNA was synthesized using ReverTra Ace qPCR RT Master Mix (TOYOBO), and quantitative PCR analysis was performed using THUNDERBIRD SYBR qPCR Mix (TOYOBO) on a StepOnePlus Real-Time PCR System (Thermo Fisher), according to the manufacturer’s instructions.

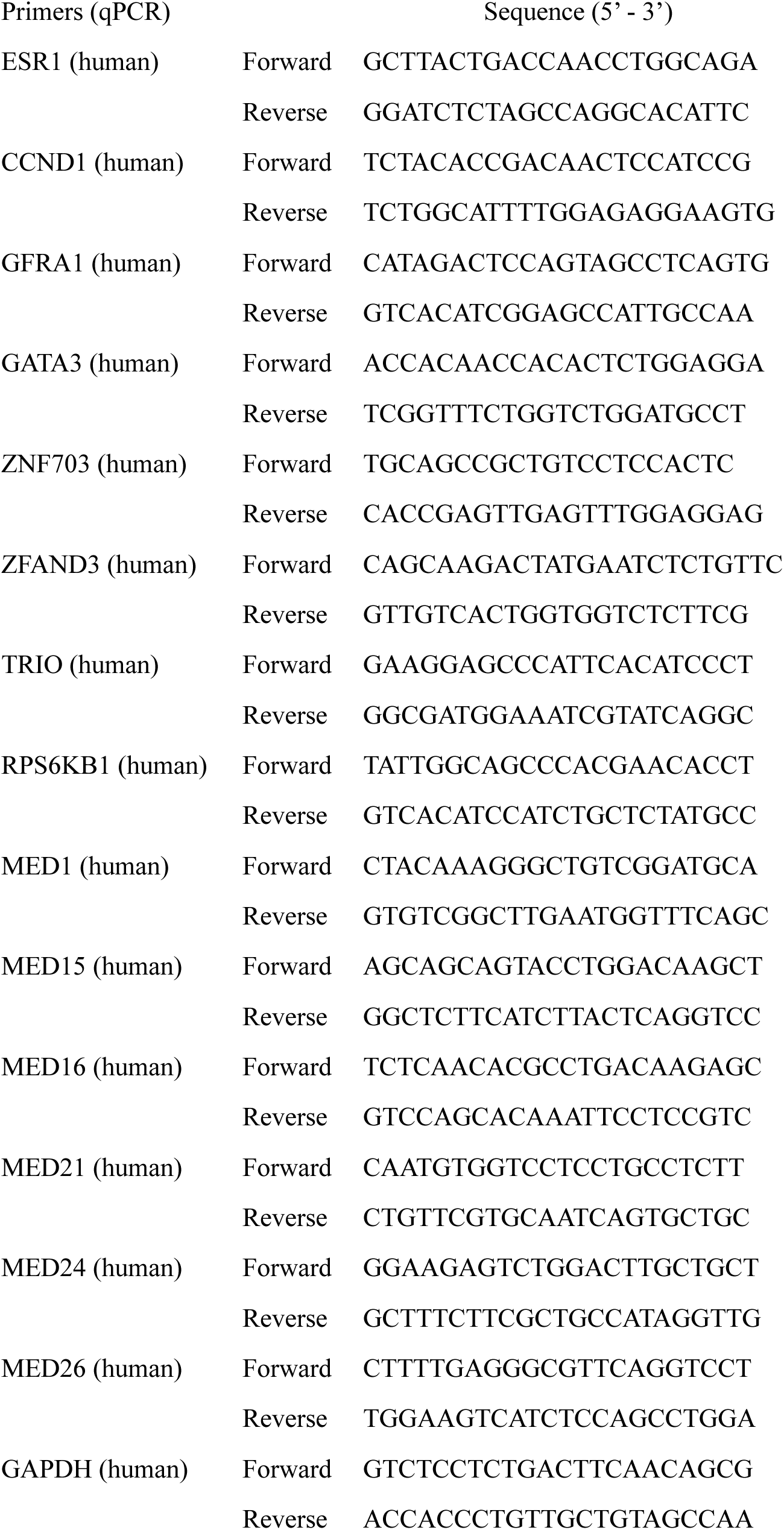
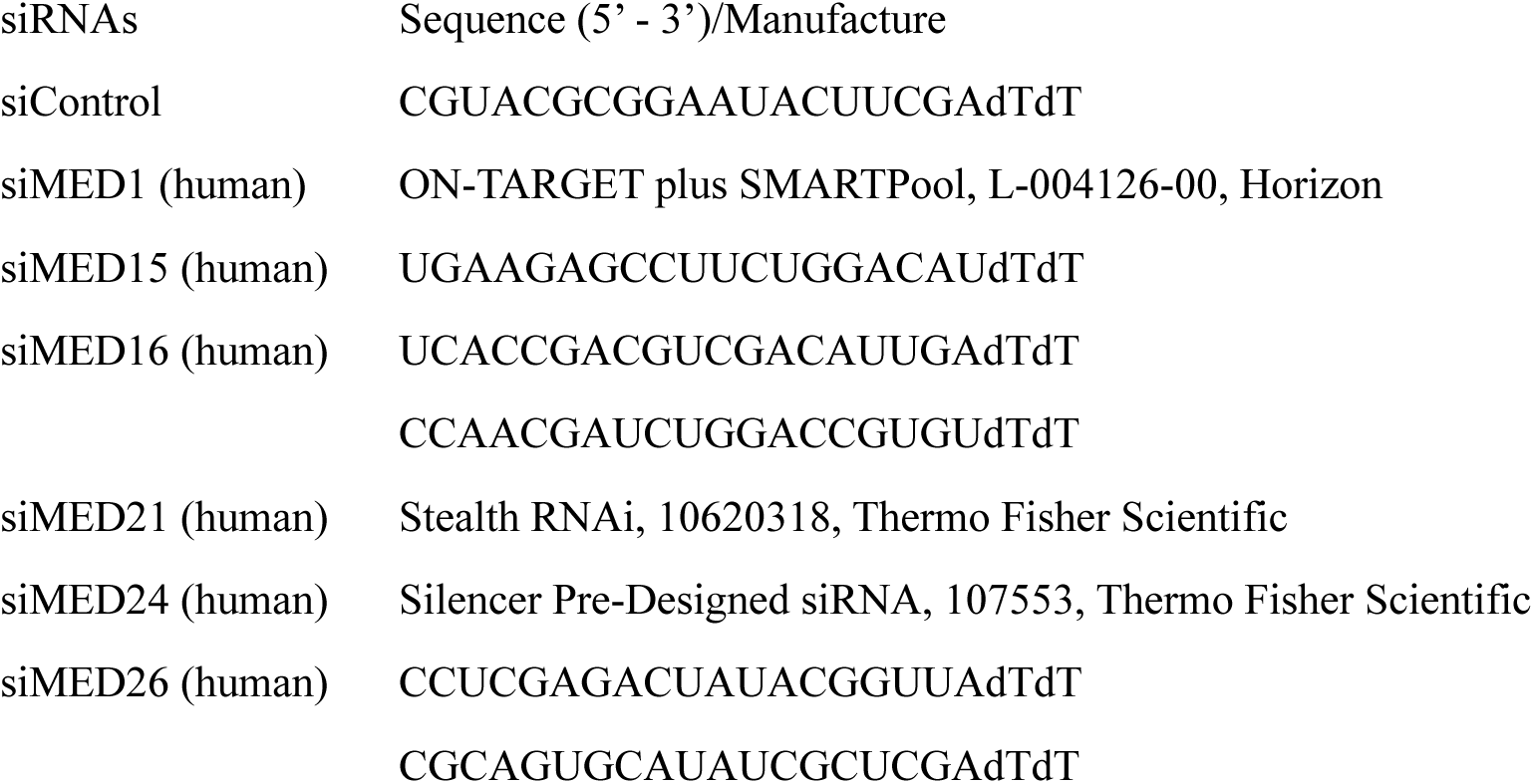

### CRISPR/Cas9-mediated dTAG (FKBP12^F36V^) knock-in cell line generation

To generate an MCF-7 cell line with a dTAG (FKBP12^F36V^) knock-in at the N-terminus of the MED24 gene, CRISPR/Cas9-mediated genome editing was performed using a crRNA/tracrRNA complex and an HDR donor template.

The guide RNA (gRNA) sequence (TGCTCAGAGTGAAATAATGAAGG) was designed using the IDT online tool and chemically synthesized by IDT as a crRNA. The crRNA was complexed with a tracrRNA and recombinant Cas9 protein (IDT) to form a ribonucleoprotein (RNP) complex before transfection. A double-stranded HDR donor template (Donor Blocks, IDT) was used for homology-directed repair. The donor template contained 200 bp homology arms flanking the knock-in site and a tandem HA tag followed by the dTAG sequence.

The inserted sequence was as follows:

TACCCCTACGACGTGCCCGACTACGCCGGCTATCCGTATGATGTCCCGGACTATGCAGGAGTGCAGGTGGAAACCATCTCCCCAGGAGACGGGCGCACCTTCCCCAAGCGCGGCCAGACCTGCGTGGTGCACTACACCGGGATGCTTGAAGATGGAAAGAAAGTTGATTCCTCCCGGGACAGAAACAAGCCCTTTAAGTTTATGCTAGGCAAGCAGGAGGTGATCCGAGGCTGGGAAGAAGGGGTTGCCCAGATGAGTGTGGGTCAGAGAGCCAAACTGACTATATCTCCAGATTATGCCTATGGTGCCACTGGGCACCCAGGCATCATCCCACCACATGCCACTCTCGTCTTCGATGTGGAGCTTCTAAAACTGGAA.

MCF-7 cells were then electroporated with the Cas9-guide RNA complex and the HDR donor DNA using the Neon Transfection System (Thermo Fisher), according to the manufacturer’s protocol. After electroporation, cells were seeded and expanded for colony formation in RPMI medium. To genotype the knock-in clones, PCR screening was conducted on 384 picked colonies, and HA-dTAG-tagged MED24 expression was confirmed by western blot.

### Cell viability assay

Cell viability was assessed using a Cell Counting Kit-8 (CCK-8; Dojindo). A total of 2,000 cells per well were seeded in a 96-well plate and cultured for 24 h. Subsequently, 500 nM dTAGV-1 or an equivalent volume of DMSO was added to the wells. At 0, 24, 48, 72, and 96 h after treatment, the CCK-8 solution was added according to the manufacturer’s instructions, and the absorbance was measured at 450 nm using a microplate reader.

### Estradiol treatment of the knock-in cells

The MCF-7 cells with the dTAG knock-in were cultured in phenol red-free medium supplemented with charcoal-stripped fetal bovine serum (FBS) for 48 h, to minimize endogenous estrogenic effects. After 45 h of culture, cells were treated with either 500 nM dTAG^V^-1 or an equivalent volume of DMSO. At 48 h, cells were further treated with or without 10 nM β-estradiol (Fujifilm). After 3 h of estradiol stimulation, total RNA was extracted using a Maxwell RSC SimplyRNA Cells Kit (Promega). The expression levels of the *GREB1* gene were analyzed by RT-qPCR, using *GAPDH* as an internal control.

### ChIP-seq data analysis

The raw sequencing reads were first trimmed to remove adapter sequences and low-quality bases, using Trim Galore. The processed reads were subsequently aligned to the human reference genome (hg38), using Bowtie2 under default settings. For samples subjected to spike-in normalization, the data were processed as previously described.^57^ The reads were aligned to both the human genome (GRCh38) and the fly genome (BDGP Release 6 + ISO1 MT/dm6), using Bowtie2. Following alignment, PCR duplicates were removed using the MarkDuplicates tool from Picard.

Normalization factors were calculated based on the number of reads mapped to the fly genome, and these factors were used to adjust the sequencing depth of human reads. To achieve normalization, downsampling of human genome-mapped reads was performed using the view function in SAMtools. BigWig files were generated at 1 bp intervals, and signal intensities were normalized to CPM (Counts Per Million reads) for standard normalization or left unnormalized for spike-in adjusted data, using deepTools. To compare differences between datasets, bigWig files representing signal differences were generated using bamCompare in deepTools. ChIP-seq signals were visualized using the Integrative Genomics Viewer (IGV), and aggregation plots were generated using computeMatrix and plotProfile in deepTools. Peak regions for each dataset were identified using MACS with a cutoff of 10^-5^. For gene-specific RNAPII peak scores, peak regions within each gene were extracted using intersectBed from BEDTools, and the total score was computed per gene. To identify enhancers e#1–5 in this study, we detected genomic regions exhibiting bidirectional transcription of enhancer RNAs (eRNAs), a hallmark of active enhancers, based on single strand CAGE-seq (ssCAGE-seq) data. To exclude promoter-associated signals, regions overlapping annotated mRNA transcription start sites were removed, and only intergenic and intronic signals with sufficient expression (≥2 counts in at least one sample) were retained. The resulting regions were extended by ±2 kbp for downstream analysis. Next, the regions where the RNAPII peaks overlapped with the bidirectional eRNA-producing regions were identified using intersectBed from BEDTools, and they were defined as the enhancers. Super-enhancer regions were identified using the ChIP-seq data of H3K27ac, H3K4me1, and input data with the findPeaks function in HOMER.

### Analysis of peak distributions relative to gene features

To localize Pol II binding peaks either on promoter, gene body, or intergenic regions (Figure 4B), a custom Python script (Data S1) was developed and executed, as below. The promoters and gene bodies were defined on actively transcribed genes with TPM (Transcripts Per Million) ≥1 in the RNA-seq data. Promoters were identified around the transcription start sites (TSS) of these genes, according to a previous report, which indicated a promoter region of roughly ±250 bp^59^. However, in this study the range was extended to ±1 kbp, to comprehensively capture the promoter regions and include potential regulatory elements in the analysis. Intergenic regions were defined as all remaining genomic regions. RNAPII peaks overlapping with promoter regions were classified as promoter-associated RNAPII peaks. Peaks that did not overlap with promoter regions were further examined for overlapping with the gene body, and these were classified as gene body peaks. Finally, peaks that did not fall within either promoters or gene bodies were categorized as intergenic peaks.

### Detection of chromatin interactions of enhancers using Hi-C data

To detect the promoters that interact with the identified enhancers, custom Python scripts (Data S2 and S3) were developed and executed. The Hi-C loop data (ENCFF797NNQ.bedpe) from the ENCODE database and the enhancer region data obtained in this study were processed using the pandas library. Chromatin loop anchors that contained enhancer regions were first extracted.

Subsequently, the analysis was performed to specifically assess the relationship between these enhancer-interacting regions and the expressed gene promoters, which were the same as those described in the previous section. For this purpose, overlapping regions were further intersected with promoter regions. Promoters that overlapped with these enhancer-associated regions through chromatin loops, where one anchor was associated with an enhancer and the other with a promoter, were classified as distal promoters. Additionally, an alternative approach was applied to detect proximal promoters, where the identified enhancers overlap with the gene promoters. To compute the chromatin interaction frequency for each enhancer, the number of overlapping Hi-C loop anchor points was counted.

### RNA-seq and GRO-seq data analysis

For both RNA-seq and GRO-seq, raw sequencing reads were first trimmed to remove adapter sequences and low-quality bases, using Trim Galore. The trimmed reads were then aligned to the human reference genome (hg38), using HISAT2 under default settings. To visualize genome-wide read distribution, bigWig files were generated using deepTools and normalized as TPM (Transcripts Per Million) reads. For the RNA-seq analysis, transcription expression levels were quantified using TPMCalculator, and an MA plot was generated to compare gene expression differences. For the gene ontology (GO) analysis, gene sets of interest were analyzed using findMotifs.pl in HOMER, which also provided a gene set enrichment analysis (GSEA) based on MSigDB.

### CAGE-seq data analysis

Sequencing data were processed using the CAGEfightR package (ver. 1.18.0) in R to detect transcription start sites (TSSs) and enhancers. Raw reads were aligned to the hg38 genome using STAR (ver. 2.5.0a). Mapped reads were converted into strand-separated bigWig files for visualization and analysis using deepTools (ver. 3.5.0). CAGE Transcription Start Sites (CTSSs) were quantified using quantifyCTSS() from CAGEfightR, excluding mitochondrial reads (chrM). Tag Clusters (TCs) were identified with quickTSSs() from CAGEfightR, and TSS expression was calculated and normalized in TPM. TSSs with low expression were removed (TPM ≥ 0.5 in at least one sample). Bidirectional clusters (BCs) were identified with quickEnhancers() and filtered for high expression (count number ≥ 2 in at least one sample). Based on the UCSC hg38 annotation, only intergenic and intronic BCs were retained as putative enhancers. Enhancer coordinates were extracted, formatted, and saved as MCF-7_enhancer.bed.

### 4C-seq data analysis

Sequenced 4C reads were processed with w4Cseq for alignment to the reference genome (hg38) and the interaction regions were identified.

### ATAC-seq data preparation

ATAC-seq data were analyzed using R. The TCGA ATAC-seq data of primary breast tumors were obtained from the National Cancer Institute Genomic Data Commons (GDC) (https://gdc.cancer.gov/about-data/publications/ATACseq-AWG), as described previously.^60^ The raw counts matrix (BRCA_raw_counts.txt) and the log2-normalized counts matrix (BRCA_log2norm.txt) were processed to generate a SummarizedExperiment object (TCGA_BRCA_ATAC_UQ_se.rds), as described below. The genomic coordinates of ATAC-seq peaks were extracted from BRCA_raw_counts.txt and stored as a GRanges object. The corresponding raw counts and log2-normalized counts were converted into matrices and stored in the counts and normcounts assays of the SummarizedExperiment object.

Clinical annotations of PAM50 subtype classifications were retrieved from the Xena Functional Genomics Explorer TCGA Hub (https://xenabrowser.net/hub/; GDC TCGA Breast Cancer (BRCA)) and GDC Clinical Biotab files. These annotations were integrated into the colData() slot of the SummarizedExperiment object.

The final filtered dataset (TCGA_BRCA_ATAC_UQ_se) containing T1 replicate samples with integrated clinical annotations was saved as an RDS file for downstream analyses.

### Identification of ATAC-seq peaks overlapping with enhancers

The genomic regions overlapping with enhancers were defined using a BED file of enhancer regions, as described in the ChIP-seq data analysis section, which was imported using import() from the rtracklayer package in R. The genomic coordinates of ATAC-seq peaks were obtained from rowRanges(), and enhancer-overlapping peaks were identified using overlapsAny() from the GenomicRanges package. Peaks overlapping multiple enhancer categories were removed to ensure mutually exclusive enhancer groups.

### Heatmap visualization of ATAC-seq signals that overlap with enhancers

To visualize ATAC-seq signal intensities in enhancer-overlapping regions, the ATAC-seq signal matrix for these regions was extracted from the normcounts assay of the SummarizedExperiment object (TCGA_BRCA_ATAC_UQ_se.rds). The ATAC-seq signal matrix was sorted in the order of Basal, LumA, LumB, HER2, Normal-Like, and NA subtypes. A heatmap was generated using pheatmap() from the pheatmap package, with sample subtype annotations incorporated.

### UMAP analysis of ATAC-seq signals overlapping with enhancers

The ATAC-seq signal matrix for enhancer regions was extracted from the normcounts assay of the SummarizedExperiment object (TCGA_BRCA_ATAC_UQ_se.rds), and only the genomic regions overlapping with the selected enhancers were included in the analysis. To reduce dimensionality, UMAP was applied using umap() from the uwot package with the following parameters: n_neighbors = 15, min_dist = 0.1, metric = "euclidean", n_components = 2. The resulting UMAP coordinates were stored in a data frame, annotated with PAM50 subtypes from the clinical metadata, and visualized using ggplot2. To determine the optimal number of clusters, the GAP statistic was computed using clusGap() from the cluster package, with K values ranging from 1 to 10 and 50 bootstrap replicates. The optimal number of clusters was determined using maxSE().

### Quantification of variability in chromatin accessibility

To assess the variability in chromatin accessibility across different enhancer categories, the variance in ATAC-seq normalized counts was computed for each peak in each enhancer region, using rowVars() from the matrixStats package. A violin plot was generated using ggplot2 to compare variance distributions, and statistical significance was computed using the wilcox.test() function.

### Analysis of the correlation between ATAC-seq signals at HD enhancers and gene expression levels

To assess the correlation between ATAC-seq signal intensities at the HD enhancers and gene expression patterns, the RNA-seq data of breast tumors were obtained from the Genomic Data Commons (GDC) using the functions GDCquery(), GDCdownload(), and GDCprepare() from the TCGAbiolinks package. For this analysis, we selected the top 1,000 most variably expressed genes across patient samples. Primary Tumor samples were selected to generate the SummarizedExperiment object. Spearman’s rank correlation was computed between the ATAC-seq signal at HD enhancers and the gene expression levels of 1,000 highly variable genes, which were selected based on the highest variance in expression across all samples, including different subtypes. The correlation matrix was generated, and hierarchical clustering was applied using the Ward.D2 method to group genes and ATAC-seq peaks based on their correlation patterns. Based on hierarchical clustering, the HD enhancers were classified into three clusters, and for each sample, the mean ATAC-seq signal intensity was computed for each cluster. The resulting correlation heatmap was visualized using ComplexHeatmap, and the hierarchical clustering results were saved for further classification of HD enhancer-associated regulatory patterns.

### eQTL filtering and intersection with enhancer regions

All analyses were performed in R using the dplyr, readr, stringr, and data.table packages, and bedtools was used for genomic interval operations (Data S4). To identify regulatory variants associated with transcriptional activity, the eQTL data from the Breast Mammary Tissue dataset (Breast_Mammary_Tissue.allpairs.txt) were analyzed. Variants were filtered to retain those with a minor allele frequency (MAF) greater than 0.05 and a nominal p-value below 0.05, ensuring the inclusion of statistically significant common eQTLs. From the variant_id field, genomic coordinates were extracted by parsing the chromosome and position information, and a BED file was generated by converting the coordinates to a 0-based format in accordance with the BED specification. This BED-formatted eQTL dataset was then intersected with the enhancer regions, using the bedtools intersect function with the -wa -wb options, allowing for the identification of overlapping genomic intervals. The intersected variants were matched back to the filtered eQTL table using chromosome, position, and variant ID as keys, enabling the recovery of full eQTL attributes for the overlapping set. The final dataset, comprising eQTLs located within enhancer regions, was saved as a tab-delimited file for downstream analysis.

## Data availability

The ChIP-seq, GRO-seq, RNA-seq, and 4C-seq datasets generated in this study have been deposited in the Gene Expression Omnibus (GEO) under the following accession numbers: GEO:GSE296103 (ChIP-seq), GEO:GSE296104 (GRO-seq), GEO:GSE296105 (RNA-seq) and GEO:GSE296106(4C-seq). CAGE-seq data were utilized for this study under accession number GEO:GSE299910. Hi-C data from the ENCODE project were utilized for this study under accession number ENCSR660LPJ (https://www.encodeproject.org). ATAC-seq data of primary breast tumors in TCGA as part of *The Chromatin Accessibility Landscape of Primary Human Cancers* (https://gdc.cancer.gov/about-data/publications/ATACseq-AWG) were used.^47^

The original code used in this research has been provided as Supplementary Data and uploaded as Data S1–S4. Any questions regarding the data and code should be addressed to the lead contact.

## Acknowledgments

This work was supported by JSPS KAKENHI Grant Numbers 20H05397, 22H04704 (to H.T.), JP24H01382, 23H00411 (to N.S.), and Japan Agency for Medical Research and Development (AMED) Grant Number JP23ama221428231 (to N.S.). H.T. is supported by the Takeda Science Foundation. N.S. is supported by Daiichi Sankyo Foundation of Life Science and the Takeda Science Foundation.

## Author contributions

H.T. and N.S. conceived the study and supervised the experiments. H.T. performed most of the experiments, except for CAGE-seq, with assistance from R.M., A.K., N.Y., and T.M. The CAGE-seq experiments were conducted by X.S., M.K., and P.C., who also contributed to data processing. A.O. and A.K. performed the data processing of 4C-seq. H.T. conducted bioinformatics analyses and analyzed clinical data, with additional contributions from K.K. and R.M. to the clinical data analysis. A.O., X.S., M.K., A.I., P.C., H.K., A.K., and Y.D. provided critical reagents, technical expertise, and valuable suggestions. H.T. and N.S. analyzed the data and wrote the manuscript with input from all co-authors.

## Online content

Supplemental information can be found online at XXX

---

**Figure S1.**
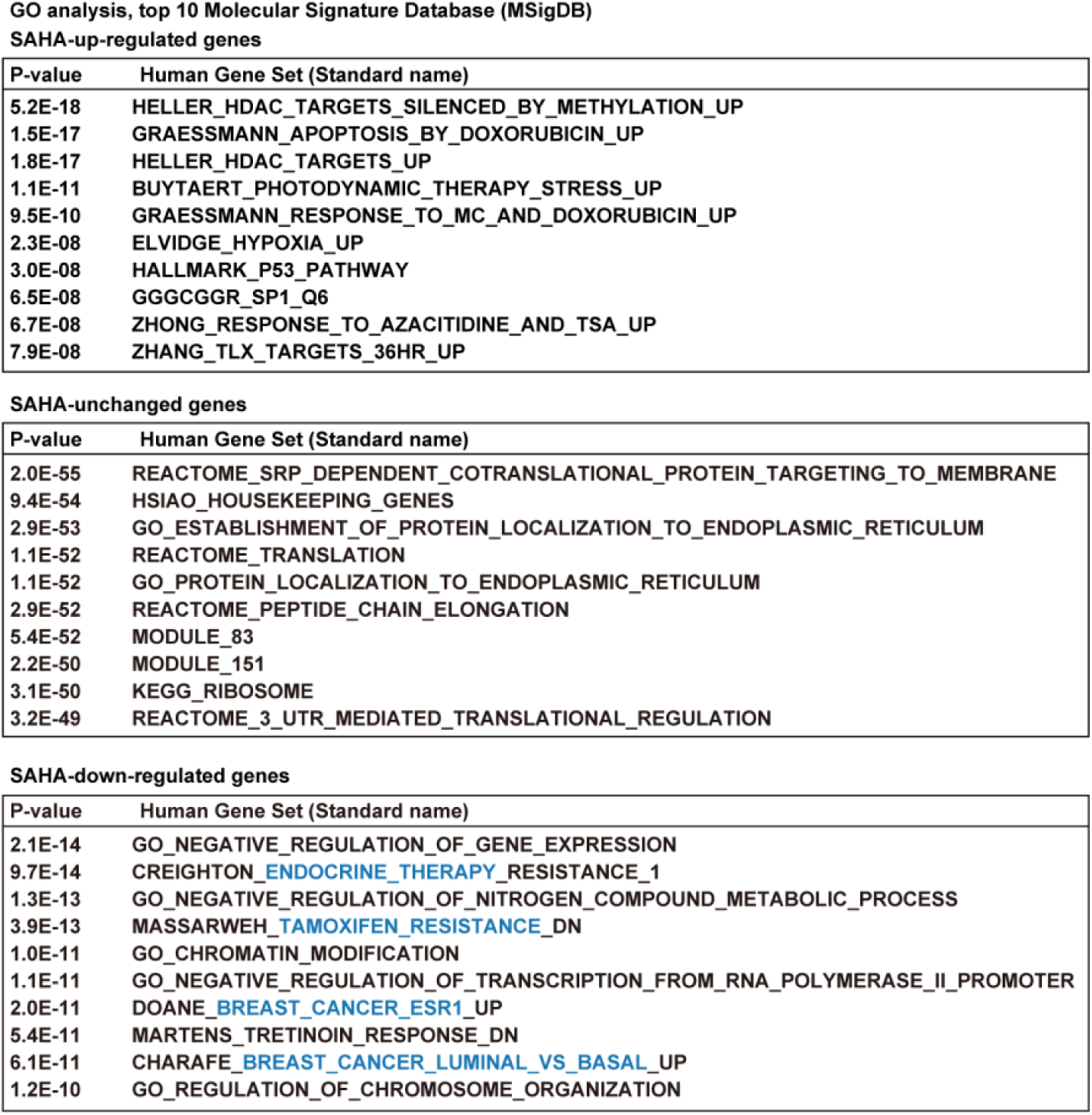
GO analysis of differentially expressed genes. The top 10 GO terms of MSigDB (GSEA) for up-regulated (upper), unchanged (middle), and down-regulated (lower) genes in MCF-7 cells treated with 2 μM SAHA for 3 h. Down-regulated genes specifically include breast cancer-related terms (blue).

**Figure S2.**
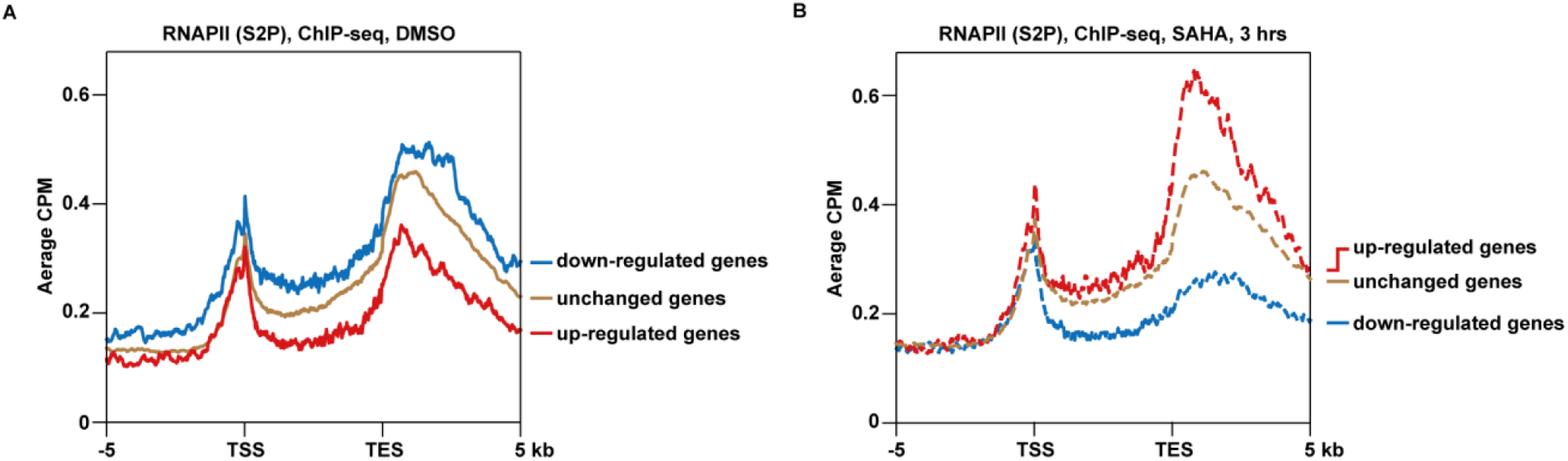
SAHA preferentially repressed transcriptionally active genes with higher RNAPII binding. (A) ChIP-seq profiling of RNAPII (S2P) in DMSO-treated MCF-7 cells. Signals for SAHA down- regulated (blue), up-regulated (red), and unchanged (brown) genes are shown. The definitions of these gene categories are based on Figure 1C. (B) The same analysis was performed after 3 h of SAHA treatment.

**Figure S3.**
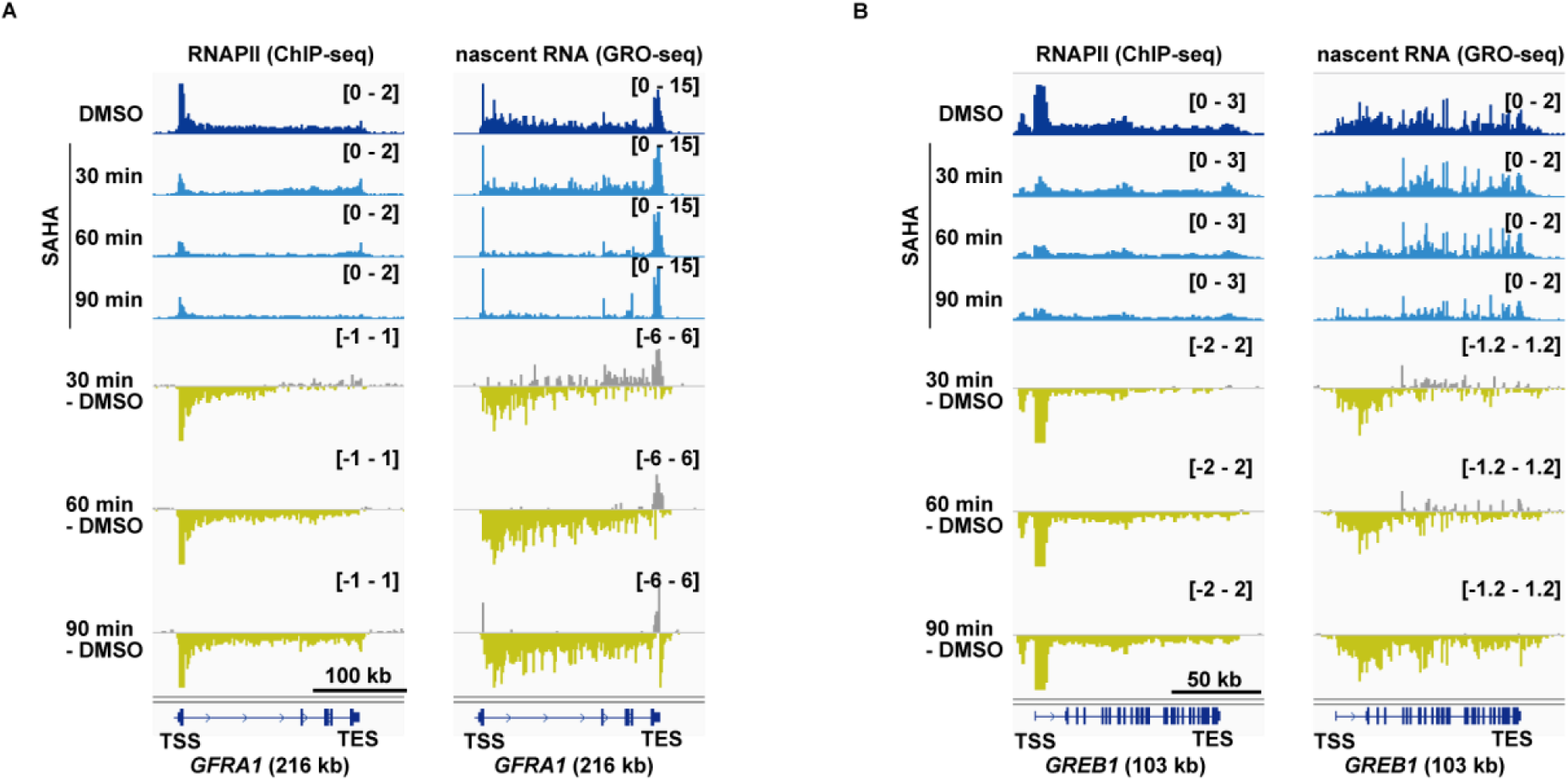
Transcriptional repression by SAHA begins with RNAPII dissociation at the 5’ end and gradually extends towards the 3’ end. (A-B) Time-course analysis of RNAPII (S5P) ChIP-seq (left) and nascent RNA GRO-seq (right) in *GFRA1* (A) and *GREB1* (B) during SAHA treatment at 30 min intervals. Transcription directions are aligned from left to right.

**Figure S4.**
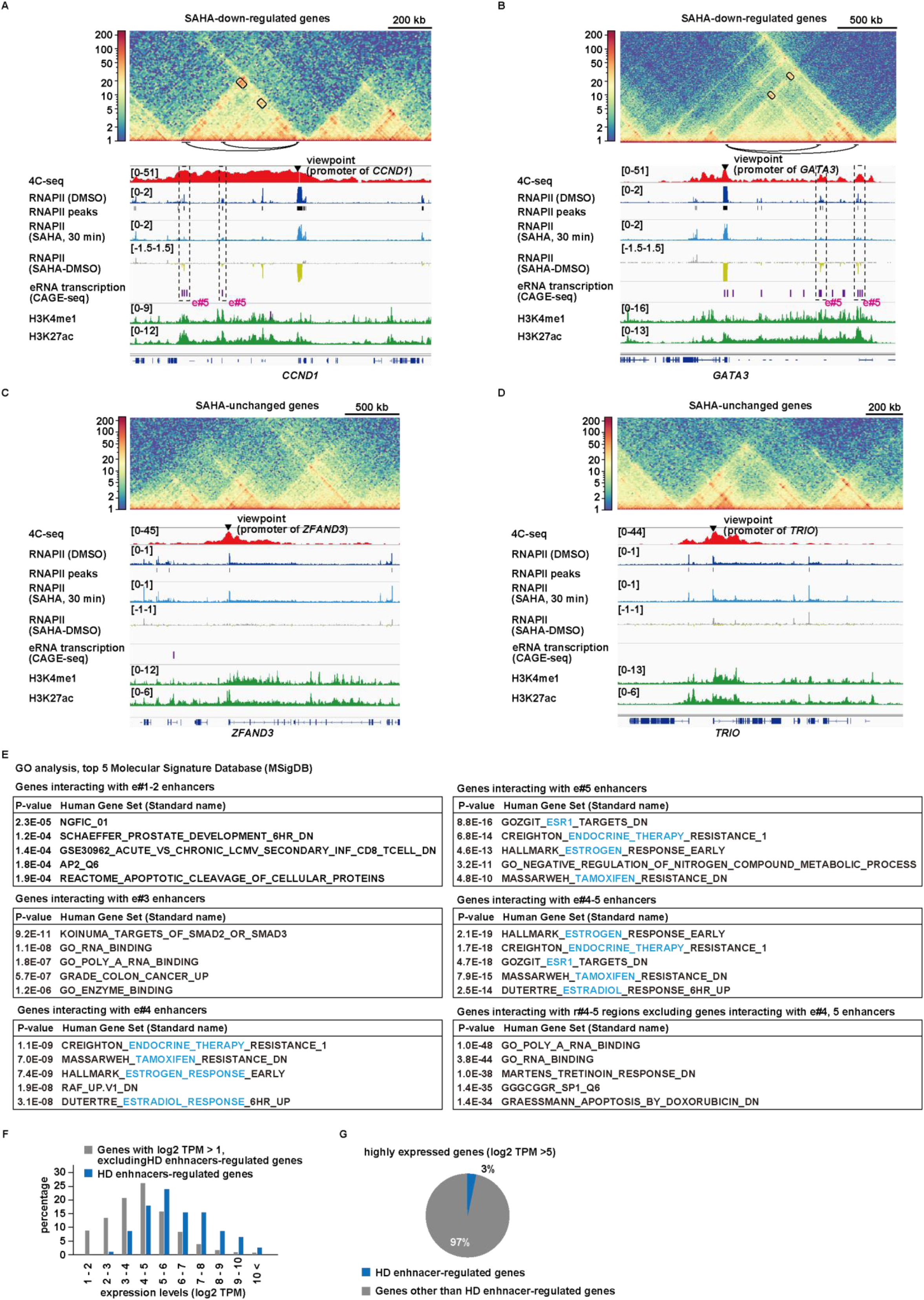
Promoters of SAHA-downregulated genes interact with enhancers from which RNAPII is dissociated by SAHA. (A-D) Alignments of Hi-C contact map, 4C-seq, RNAPII (S5P) binding, and enhancer markers (eRNA transcription) for the SAHA down-regulated genes (A and B) and SAHA-unchanged genes (C and D). RNAPII (S5P) ChIP-seq profiles are derived from the dataset used in Figure 3A. The dashed squares highlight the regions where the RNAPII binding decreases with the SAHA treatment, the enhancer markers are located, and the promoters of the indicated genes interact (A and B). Unlike SAHA-downregulated genes, SAHA-unchanged genes showed no promoter-enhancer interactions (C and D). (E) GO analysis of genes interacting with each enhancer (e#1-5). The top 5 terms from MSigDB are shown. Genes interacting with HD enhancers (e#4-5) are associated with breast cancer and highlighted in blue. (F) Genes interacting with HD enhancers (e#4-5) have a higher proportion of highly expressed genes as compared to all protein-coding genes with detectable expression. (G) Only 3% of highly expressed genes (log2 TPM > 5) are regulated (contacted) by HD enhancers.

**Figure S5.**
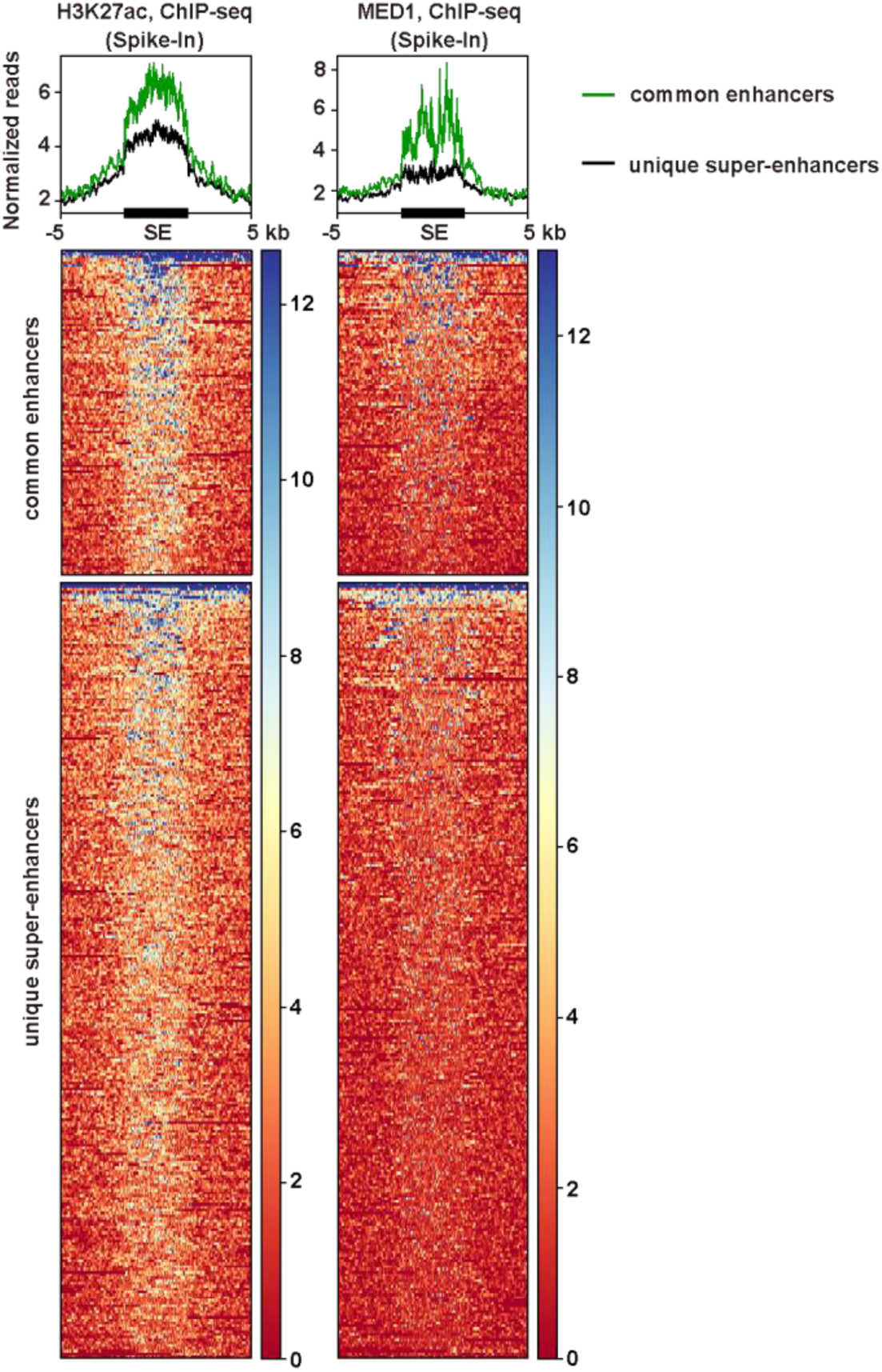
Super-enhancers containing HD enhancers show higher levels of H3K27ac and MED1 binding. Aggregation plots and heatmaps, showing H3K27ac (left) and MED1 (right) binding at common enhancers and unique super-enhancers. These enhancers are defined in Figure 5. All ChIP-seq data were normalized using spike-in controls.

**Figure S6.**
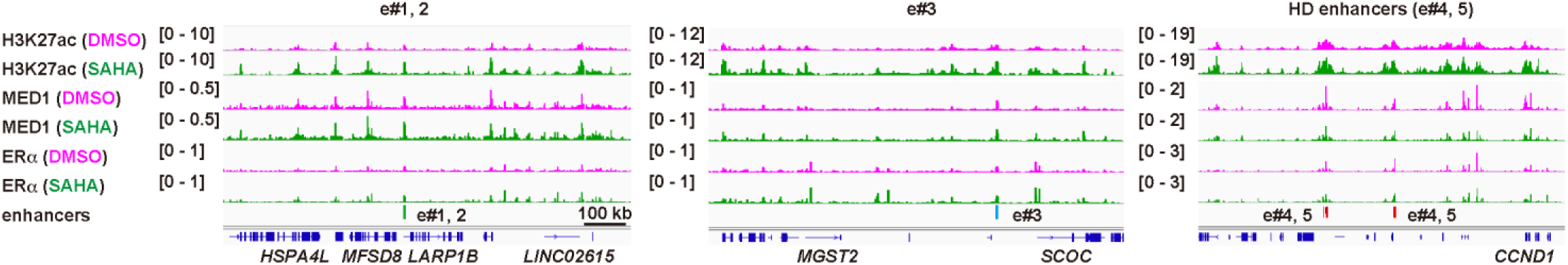
MED1 is specifically dissociated from HD enhancers upon SAHA treatment. Representative IGV images of ChIP-seq data around the enhancers defined in Figure 4C. Regions around e#1, 2 (left), e#3 (center), and HD enhancers (e#4,5, right) are shown. Results are from MCF-7 cells treated with DMSO (magenta) and SAHA (green) for 30 min.

**Figure S7.**
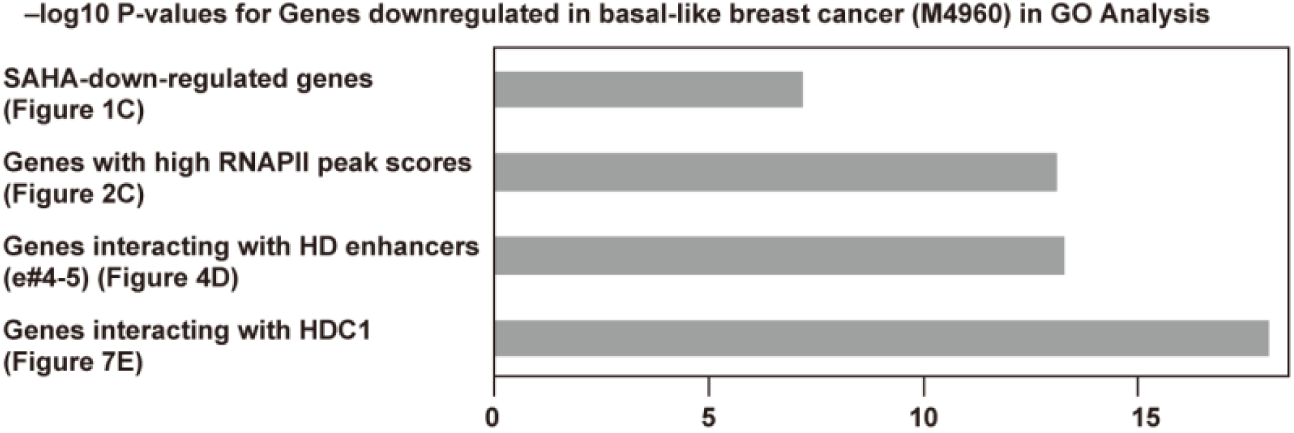
HDC1 enhancers regulate genes enriched for luminal breast cancer-associated functions. GO analysis showing that the genes downregulated in basal-like breast cancer are enriched in SAHA-down-regulated genes (shown in Figure 1C), genes with high RNAPII peak scores (shown in Figure 2C), and genes interacting with HD and HDC1 enhancers (shown in Figures 4D and 7E, respectively), in that order.

## References

1. Dawson, S., Rueda, O. M., Aparicio, S. & Caldas, C. A new genome-driven integrated classification of breast cancer and its implications. EMBO J. 32, 617–628 (2013).

2. Parker, J. S. et al. Supervised Risk Predictor of Breast Cancer Based on Intrinsic Subtypes. J. Clin. Oncol. 27, 1160–1167 (2009).

3. Yang, J. et al. Enhancer reprogramming: critical roles in cancer and promising therapeutic strategies. Cell Death Discov. 11, 84 (2025).

4. Patten, D. K. et al. Enhancer mapping uncovers phenotypic heterogeneity and evolution in patients with luminal breast cancer. Nat Med 24, 1469–1480 (2018).

5. Fachal, L. et al. Fine-mapping of 150 breast cancer risk regions identifies 191 likely target genes. Nat. Genet. 52, 56–73 (2020).

6. Nasser, J. et al. Genome-wide enhancer maps link risk variants to disease genes. Nature 593, 238–243 (2021).

7. González, A. & Paul, P. Pleiotropic expression quantitative trait loci are enriched in enhancers and transcription factor binding sites and impact more genes. Comput. Struct. Biotechnol. J. 23, 4260–4270 (2024).

8. Creyghton, M. P. et al. Histone H3K27ac separates active from poised enhancers and predicts developmental state. Proc. Natl. Acad. Sci. 107, 21931–21936 (2010).

9. Heintzman, N. D. et al. Histone modifications at human enhancers reflect global cell-type- specific gene expression. Nature 459, 108–112 (2009).

10. Heintzman, N. D. et al. Distinct and predictive chromatin signatures of transcriptional promoters and enhancers in the human genome. Nat. Genet. 39, 311–318 (2007).

11. Weintraub, H. A dominant role for DNA secondary structure in forming hypersensitive structures in chromatin. Cell 32, 1191–1203 (1983).

12. Khoury, G. & Gruss, P. Enhancer elements. Cell 33, 313–314 (1983).

13. Thurman, R. E. et al. The accessible chromatin landscape of the human genome. Nature 489, 75–82 (2012).

14. Kim, T.-K. et al. Widespread transcription at neuronal activity-regulated enhancers. Nature 465, 182–187 (2010).

15. Oguchi, A. et al. An atlas of transcribed enhancers across helper T cell diversity for decoding human diseases. Science 385, eadd8394 (2024).

16. Andersson, R. et al. An atlas of active enhancers across human cell types and tissues. Nature 507, 455–461 (2014).

17. Melgar, M. F., Collins, F. S. & Sethupathy, P. Discovery of active enhancers through bidirectional expression of short transcripts. Genome Biol. 12, R113–R113 (2011).

18. Consortium, T. E. P. et al. Expanded encyclopaedias of DNA elements in the human and mouse genomes. Nature 583, 699–710 (2020).

19. Daugherty, A. C. et al. Chromatin accessibility dynamics reveal novel functional enhancers in C. elegans. Genome Res. 27, 2096–2107 (2017).

20. Zu, S. et al. Single-cell analysis of chromatin accessibility in the adult mouse brain. Nature 624, 378–389 (2023).

21. Whalen, S., Truty, R. M. & Pollard, K. S. Enhancer–promoter interactions are encoded by complex genomic signatures on looping chromatin. Nat. Genet. 48, 488–496 (2016).

22. Schoenfelder, S. & Fraser, P. Long-range enhancer–promoter contacts in gene expression control. Nat. Rev. Genet. 20, 437–455 (2019).

23. Tolhuis, B., Palstra, R.-J., Splinter, E., Grosveld, F. & Laat, W. de. Looping and Interaction between Hypersensitive Sites in the Active β-globin Locus. Mol. Cell 10, 1453–1465 (2002).

24. Sanyal, A., Lajoie, B., Jain, G. & Dekker, J. The long-range interaction landscape of gene promoters. Nature 489, 109–113 (2012).

25. Rengachari, S., Schilbach, S., Aibara, S., Dienemann, C. & Cramer, P. Structure of the human Mediator–RNA polymerase II pre-initiation complex. Nature 594, 129–133 (2021).

26. Allen, B. L. & Taatjes, D. J. The Mediator complex: a central integrator of transcription. Nat. Rev. Mol. Cell Biol. 16, 155–166 (2015).

27. Richter, W. F., Nayak, S., Iwasa, J. & Taatjes, D. J. The Mediator complex as a master regulator of transcription by RNA polymerase II. Nat. Rev. Mol. Cell Biol. 23, 732–749 (2022).

28. Jaeger, M. G. et al. Selective Mediator dependence of cell-type-specifying transcription. Nat. Genet. 52, 719–727 (2020).

29. Bhagwat, A. S. et al. BET Bromodomain Inhibition Releases the Mediator Complex from Select cis-Regulatory Elements. Cell Rep. 15, 519–530 (2016).

30. Whyte, W. A. et al. Master Transcription Factors and Mediator Establish Super-Enhancers at Key Cell Identity Genes. Cell 153, 307–319 (2013).

31. Wang, Z. et al. Genome-wide Mapping of HATs and HDACs Reveals Distinct Functions in Active and Inactive Genes. Cell 138, 1019–1031 (2009).

32. LaBonte, M. J. et al. The Dual EGFR/HER2 Inhibitor Lapatinib Synergistically Enhances the Antitumor Activity of the Histone Deacetylase Inhibitor Panobinostat in Colorectal Cancer Models. Cancer Res. 71, 3635–3648 (2011).

33. Scott, G. K. et al. Destabilization of ERBB2 Transcripts by Targeting 3′ Untranslated Region Messenger RNA Associated HuR and Histone Deacetylase-6. Mol. Cancer Res. 6, 1250–1258 (2008).

34. Kelly, R. D. W. et al. Histone deacetylases maintain expression of the pluripotent gene network via recruitment of RNA polymerase II to coding and noncoding loci. Genome Res. 34, 34–46 (2024).

35. Tao, R. et al. Deacetylase inhibition promotes the generation and function of regulatory T cells. Nat. Med. 13, 1299–1307 (2007).

36. Grozinger, C. M. & Schreiber, S. L. Deacetylase Enzymes Biological Functions and the Use of Small-Molecule Inhibitors. Chem. Biol. 9, 3–16 (2002).

37. Gryder, B. E. et al. Chemical genomics reveals histone deacetylases are required for core regulatory transcription. Nat Commun 10, 3004 (2019).

38. Slaughter, M. J. et al. HDAC inhibition results in widespread alteration of the histone acetylation landscape and BRD4 targeting to gene bodies. Cell Reports 34, 108638 (2021).

39. Lopes, R. et al. Systematic dissection of transcriptional regulatory networks by genome-scale and single-cell CRISPR screens. Sci. Adv. 7, (2021).

40. Muniz, L., Nicolas, E. & Trouche, D. RNA polymerase II speed: a key player in controlling and adapting transcriptome composition. Embo J 40, e105740 (2021).

41. Bhakta, S. et al. An Anti-GDNF Family Receptor Alpha 1 (GFRA1) Antibody-Drug Conjugate for the Treatment of Hormone Receptor-Positive Breast Cancer. Mol. cancer Ther. 17, 638–649 (2017).

42. Zhang, Y. et al. Chromatin connectivity maps reveal dynamic promoter–enhancer long-range associations. Nature 504, 306–310 (2013).

43. Deng, W. et al. Controlling Long-Range Genomic Interactions at a Native Locus by Targeted Tethering of a Looping Factor. Cell 149, 1233–1244 (2012).

44. Hnisz, D. et al. Super-Enhancers in the Control of Cell Identity and Disease. Cell 155, 934–947 (2013).

45. Pott, S. & Lieb, J. D. What are super-enhancers? Nat. Genet. 47, 8–12 (2015).

46. Lovén, J. et al. Selective Inhibition of Tumor Oncogenes by Disruption of Super-Enhancers. Cell 153, 320–334 (2013).

47. Corces, M. R. et al. The chromatin accessibility landscape of primary human cancers. Science 362, (2018).

48. Tishchenko, I., Milioli, H. H., Riveros, C. & Moscato, P. Extensive Transcriptomic and Genomic Analysis Provides New Insights about Luminal Breast Cancers. PLoS ONE 11, e0158259 (2016).

49. Ciriello, G. et al. The molecular diversity of Luminal A breast tumors. Breast Cancer Res. Treat. 141, 409–420 (2013).

50. Leidescher, S. et al. Spatial organization of transcribed eukaryotic genes. Nat Cell Biol 24, 327– 339 (2022).

51. Tantale, K. et al. A single-molecule view of transcription reveals convoys of RNA polymerases and multi-scale bursting. Nat. Commun. 7, 12248 (2016).

52. Henikoff, S. et al. RNA polymerase II at histone genes predicts outcome in human cancer. Science 387, 737–743 (2025).

53. Métivier, R. et al. Estrogen Receptor-α Directs Ordered, Cyclical, and Combinatorial Recruitment of Cofactors on a Natural Target Promoter. Cell 115, 751–763 (2003).

54. Lin, R. et al. MED1 IDR acetylation reorganizes the transcription preinitiation complex, rewires 3D chromatin interactions and reprograms gene expression. bioRxiv 2024.03.18.585606 (2024) doi:10.1101/2024.03.18.585606.

55. Hasegawa, N. et al. Mediator Subunits MED1 and MED24 Cooperatively Contribute to Pubertal Mammary Gland Development and Growth of Breast Carcinoma Cells. Mol. Cell. Biol. 32, 1483–1495 (2012).

56. Sato, S. et al. Role for the MED21-MED7 Hinge in Assembly of the Mediator-RNA Polymerase II Holoenzyme*. J. Biol. Chem. 291, 26886–26898 (2016).

57. Tachiwana, H. et al. Chromatin structure-dependent histone incorporation revealed by a genome-wide deposition assay. Elife 10, e66290 (2021).

58. Barbieri, E. et al. Rapid and Scalable Profiling of Nascent RNA with fastGRO. Cell Reports 33, 108373 (2020).

59. Butler, J. E. F. & Kadonaga, J. T. The RNA polymerase II core promoter: a key component in the regulation of gene expression. Genes Dev. 16, 2583–2592 (2002).

60. Kumegawa, K. et al. Chromatin profile-based identification of a novel ER-positive breast cancer subgroup with reduced ER-responsive element accessibility. Br. J. Cancer 128, 1208–1222 (2023).

